# Single Cell RNA Sequencing Analysis of Human Melanoma Reveals A Distinct Prognostic Myeloid Cell

**DOI:** 10.1101/2025.03.05.633739

**Authors:** M. Usman Ahmad, Lyong Heo, Ramesh V. Nair, Saurabh Sharma, Emanual Maverakis, Amanda R. Kirane

## Abstract

Myeloid-derived cells and tumor-associated macrophages (TAMs) represent an emerging field in melanoma research. Single-cell sequencing has enhanced our understanding of interactions within the tumor microenvironment (TME). Although there is no current consensus, several solid-tumor associated TAM subtypes have been identified by single-cell RNA sequencing (scRNA-seq). Due to the lack of agreement across models and tumor types, we aimed to characterize the myeloid sub-compartment in melanoma by analyzing compiled public scRNA-seq datasets. In this study, we identified six myeloid cell clusters, designated M_C1 through M_C6, with distinct gene expression profiles indicating differences in structure and function. The frequency of the M_C1 subtype was found to be predictive of overall survival (OS) in brain metastatic melanoma. This finding was externally validated by deconvolution of publicly available bulk RNA sequencing data from a clinical trial of advanced melanoma patients treated with nivolumab, as well as data from The Cancer Genome Atlas (TCGA) for melanoma stages I–IV. The M_C1 cell type appears to be a hybrid of dendritic cells and macrophages, demonstrating prognostic relevance across all melanoma stages and predictive potential for immunotherapy response. These data suggest that M_C1 may have utility as a prognostic biomarker in melanoma and as a target for future cell-based therapies.

## Introduction

While immunotherapy has revolutionized the management of advanced melanoma, modern therapeutic selection has become increasingly complex with both the adoption of neoadjuvant approaches and the recent approvals of multiple systemic agents^1^. Despite significant improvements for some patients, overall response remains unpredictable, with response rates of 10-15% for ipilimumab, 33-44% for pembrolizumab/nivolumab, 58% for ipilimumab + nivolumab, and a 48% progression free survival nivolumab + relatlimab^1^. Additionally, both inherent or acquired resistance remain challenges, such as the average resistance to dabrafenib within 1 year^1^. Our group has specifically aimed to dissect the role of myeloid cells and tumor associated macrophages (TAMs) in suppressing immune responses, contributing to anti-PD-1, and other immunotherapy failures.

Immune checkpoint inhibitors such as ipilimumab, nivolumab, and pembrolizumab target T cell related mechanisms related to PD-1/PD-L1 and CTLA-4^2,3^. Understanding of these drugs and their mechanisms within the tumor-immune microenvironment (TiME) have emphasized T cell related mechanisms, however, emerging concepts include the microbiome and checkpoint activity across other immune cell subtypes, such as the critical role of PD-L1/PD-1 signaling across TAMs^4^. Improved resolution of cells from within tumor samples using single-cell sequencing technology in melanoma has improved our understanding of cellular plasticity, T, and NK cell functional states, and the interplay of cells in tumor mediated immunosuppression^5^.

The myeloid compartment includes a variety of cell types including but not limited to neutrophils, eosinophils, basophils, monocytes, macrophages, and dendritic cells^6,7^. The emerging role of tumor associated macrophages in the tumor ecosystems and immunotherapy responses is provocative, however understanding of these roles has been complicated by the plasticity of macrophages subtype differentiation, with paradoxical functions existing based upon the exact signals in the biological environment in both native tissues and disease states^8,9^. Recently, there has been a paradigm shift in defining macrophages in the tumor immune microenvironment from a model based on “M1,” or inflammatory macrophages, and “M2,” immunosuppressive macrophages, into several subtypes. Attempts to design novel classification systems based on function have been inconsistent due to heterogeneity in multiple phenotypes across solid tumors^10–12^. Previously, analysis of scRNA sequencing data in melanoma has revealed significant overall survival differences with dendritic cell (DC) subpopulations^18^ and increased M1 type macrophages and lower immune checkpoint expression on CD8 T cells^19^. Overall, it appears that there is a possibility of specific myeloid cell sub-types unique to a particular solid tumor but a lack of consensus of myeloid cell sub-types in human melanoma. Thus, we undertook analysis of a comprehensive scRNA sequencing data set using machine learning techniques with the objective to define myeloid signatures conserved across human melanoma to define subtype populations of critical relevance by biodistribution, tumor stage or recurrence, and impacting immunotherapeutic responses.

## Results

### Cell Types in early, advanced, and metastatic Melanoma & Interaction

A total of 123 samples were included in the data set for scRNA sequencing with a total of 102,971 single cells. A total of 45 cell types including tumor cells, fibroblasts, endothelial cells, NK cells, CD3 T cells, CD4 T cells, CD8 T cells, B cells, and myeloid cells were found after clustering as depicted in **Figure 1a**. Gene expression of canonical lineage markers were evaluated to broadly classify cells into tumor, T cells, NK cells, B cells, and myeloid cells as shown in **Figure 1c**. When evaluating overall expression of cell types by tissue type there were significant differences in T cells, B cells, NK cells, extracellular matrix cells (ECM), and tumor cells as shown in **Figure 1b**. However, myeloid cell expression was not significantly different by tissue types. Individual cell types within these clusters were evaluated for differences in expression between tissues types with significant differences between groups in **Supplement 1**.

**Figure 1.**
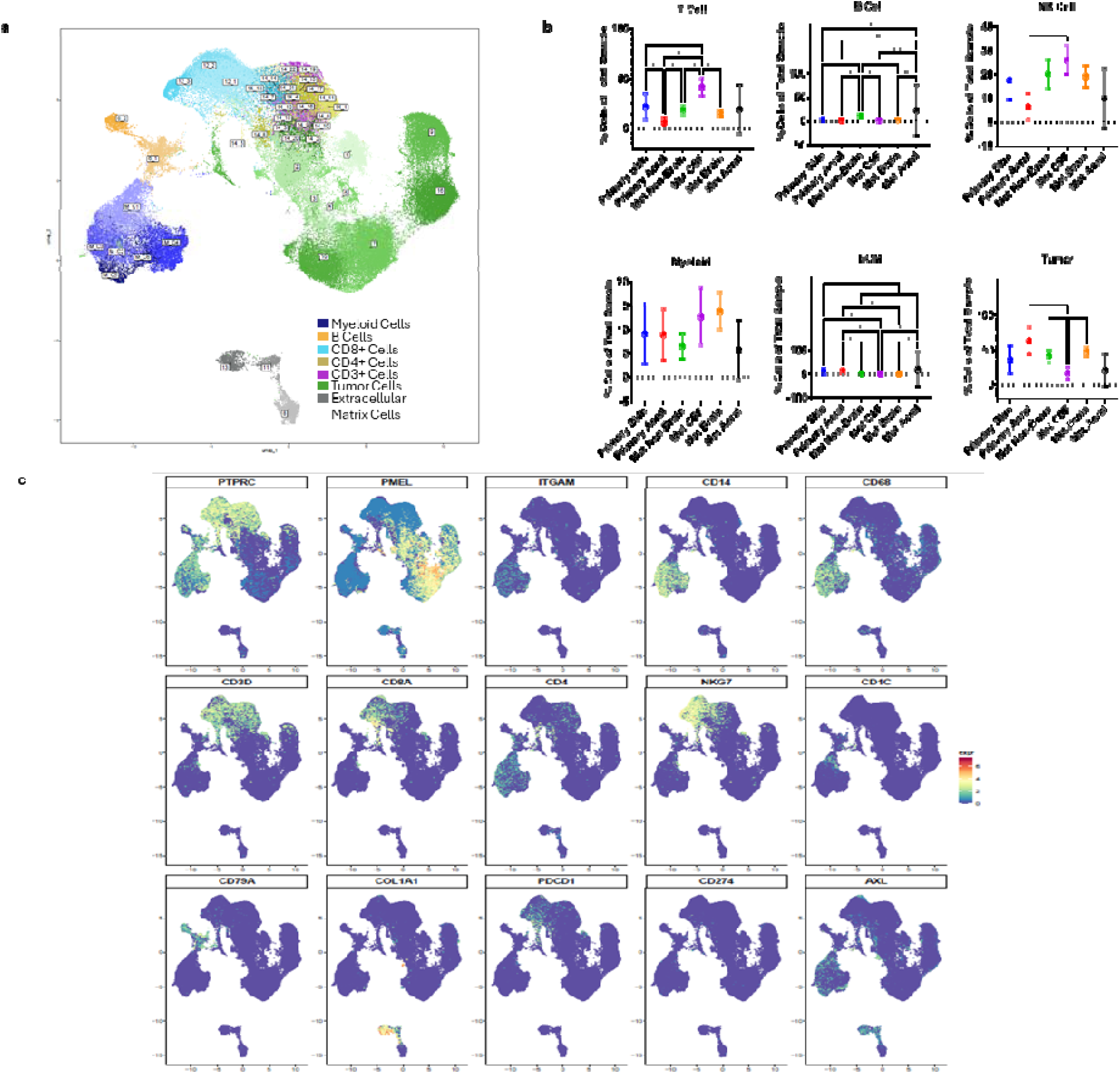
Overall Results of Clustering on single-cell RNA sequencing of Melanoma Patients. **a,** Overall clustering and sub-clustering results using scCESS depicted on two-coordinate feature map using UMAP **b,** Overall, frequency of cell type compared to all sequenced cells by tissue source analyzed for between group differences using GraphPad Prism using one way ANOVA with Tukey’s multiple comparisons testing; * p < 0.05, ** p < 0.01, *** p <0.001, **** p <0.0001. **c,** Gene expression by cluster with selected genes as log2FC value on heatmap. **d,** Heatmap of relative gene expression of selected genes on two coordinate feature UMAP.

#### Tumor Cells

13 clusters were identified with increased expression of tumor related markers including PMEL (**Figure 1a and 1c**). Cluster 3 cells were elevated in primary acral melanoma but no other groups with significant differences on ANOVA (**Supplement 1**). When evaluating this cluster by a cut-off log2FC of 2.5 with 80% expression and a 30% change in expression compared to other clusters, these cells appear to be significantly elevated in BRAF for gene expression, average Log2FC 2.98, p=0.00. Other tumor clusters with increased BRAF expression include 5, 4, and 2. Cluster 16 cells were significantly elevated in brain metastases when compared to cerebrospinal fluid metastases, non-brain metastases, and primary skin melanoma (**Supplement 1**). These cells highly express NRN1, STXBP1, VAT1, MLANA, BIRC7, and S100A13 which supports brain specific melanoma tumor cells. Clusters increased in expression in primary vs metastatic melanoma include 14_2, with increased expression CTAG2, and 14_20, with increased expression CTNNA2 (**Supplement 1**).

#### Myeloid Cells

6 clusters express ITGAM and are of myeloid origin (**Figure 1a and 1c**). Cluster M_C1 expresses ITGAX, CD14, FCGR3A, CD68, CD163, MRC1, CSF1R, PECAM1, TREM2 and uniquely FLT3. FLT3 suggests M_C1 may have features of a hybrid dendritic cell precursor and macrophage type with no significant differences by tissue type (**Supplement 1**). M_C2 has a similar profile, but does not express FLT3 and does express MARCO, pointing to an inflammatory macrophage specific differentiation with no significant differences by tissue type (**Supplement 1**). M_C3 does not express FLT3 or MARCO and has a similar profile and appears to be another inflammatory macrophage subtype with no significant differences by tissue type (**Supplement 1**). M_C6 demonstrates similar profile to M_C3 without TREM2 expression, pointing to regulatory macrophage differentiation and was increased in brain vs non-brain metastases (**Supplement 1**). M_C4 expresses SELL and does not express MRC, pointing to monocytic or inflammatory differentiation with increased expression in brain vs non-brain metastases (**Supplement 1**). M_C5 expresses similar markers to other myeloid clusters, but uniquely expresses CX3CR1 and P2RY12, with no significant differences by tissue type despite association with microglial differentiation (**Supplement 1**).

#### CD8 Expressing Cells

6 clusters express CD8A or CD8B with CD3 genes (**Figure 1a and 1c**). Clusters 12_1, 12_2, and 12_3 uniquely overexpress NKG7, GZMB, and GZMH, possibly pointing to NKT cells more polarized to NK type cells. Cluster 12_1 has high expression of CCL5 (log2FC 3.56, p=0.00) and NKG7 (log2FC 3.53, p=0.00) pointing to an activated and cytotoxic phenotype. Cluster 12_2 has high expression of GNLY (log2FC 7.03, p=0.00) and NKG7 (log2FC 3.07, p=0.00) also showing activation markers. Cluster 12_3 highly expresses CCL4, NKG7, CCL5, CST7, and GZMA complementing activation and cytotoxicity markers although CST7 may be paradoxical and CCL4 may assist in recruitment of macrophages and dendritic cells. Cluster 14_4 highly expressed exhaustion signatures IL7R, CD52, PTPRC, PRDM1 and S100A4. Cluster 14_7 expresses increased PDCD1, log2FC 1.64, p=2.37E-17, and Cluster 14_14 also expresses PDCD1 and S100A4. Clusters 14_4 and 14_14 are both more highly expressed in CSF metastases compared to other metastatic sites of disease pointing toward loss of T cell cytotoxicity in the most malignant patient sub-type (**Supplement 1**).

#### CD4 Expressing Cells

11 clusters express CD4 with CD3 genes, indicating a multitude of potential cell subsets (**Figure 1a and 1c**). Cluster 14_22 (FOXP3, IL2RA, ICOS, CTLA-4) and Cluster 14_5 (FOXP3, ICOS, CTLA-4, CCR6) appear to be two different regulatory CD4 (Treg) cells. There appear to be no significant differences in expression by total sample in 14_5 by tissue, however, 14_22 appears to be significantly elevated in CSF metastases compared to brain metastases and primary acral melanoma (**Supplement 1**). Cluster 14_13 is upregulated for PDCD1 and is significantly elevated in metastatic melanoma compared to brain metastases and primary acral melanoma (**Supplement 1**). Cluster 14_15 (IL4R high) was without differences between tissue types. Cluster 14_12 is singularly upregulated for CCR4 with increased expression in brain metastases compared to CSF metastases, non-brain metastases, primary acral, and primary skin melanoma (**Supplement 1**). Clusters 14_1 and 14_11 are upregulated for GATA3 and appear to increased in non-brain metastases compared to primary acral melanoma (Cluster 14_1) and increased compared to brain metastases, CSF metastases, primary acral, and primary melanoma (14_11) (**Supplement 1**). Cluster 14_21 appears to be uniquely upregulated for BCL-6 signifying importance as a T follicular helper cell without differences between tissues (**Supplement 1**). Clusters 14_17, 14_6, and 14_8 appears to have characteristics of early undifferentiated CD4 T cells.

#### CD3 Expressing Cells

Four clusters express CD3D, CD3E, and CD3G without CD8 or CD4 genes (**Figure 1a and 1c**). Clusters 14_10 and 14_18 are increased in metastatic acral compared to all other tissue types (**Supplement 1**). Clusters 14_16 and 14_19 are increased in CSF metastases compared to all other tissue types (**Supplement 1**).

#### Extracellular Matrix (ECM) Cells

Three clusters represent fibroblasts and endothelial cells (**Figure 1a and 1c**). Cluster 8 highly expressed endothelial cell markers CD34 and PECAM-1 and does not differ by tissue type (**Supplement 1**). Clusters 11 and 13 are both elevated in expression of COL1A1 a fibroblast marker. Cluster 11 is increased in acral metastases compared to brain metastases and non-brain metastases while Cluster 13 is not different by tissue type (**Supplement 1**).

#### B Cells

2 clusters highly express B cell associated lineage genes CD79A and CD79B (**Figure 1a and 1c**). Cluster 6_1 is increased in acral metastases compared to brain metastases, primary acral, and primary melanoma (**Supplement 1**). In addition, 6_1 is increased in non-brain metastases compared to CSF metastases, primary acral, primary melanoma, and brain metastases (**Supplement 1**).

#### Cell Chat Analysis

Myeloid clusters had outgoing upregulation of LAIR1 and APRIL signaling which have paradoxical immune effects and incoming LAIR1, CD39, and CD6 (**Figure 2a and Figure 2b**). This finding of a highly inflammatory state of myeloid cell clusters with paradoxical regulation signals underscores the complexities of myeloid functions that have rendered “M1” and “M2” classification systems inadequate to comprehensively describe cellular functions in tumors. For tumor cells, outgoing signaling patterns varied by tumor cell, however, incoming signals appeared to be highly upregulated in pattern 2 by the VISTA pathway known for downregulating immune cell function (**Figure 2b**). Outgoing signals for NKT and CD8 T cell related cell clusters included IFN-II and incoming signals CLEC and MHC-I (**Figure 2a and Figure 2b**). Other T cells overlapped with these outgoing and incoming signals (**Figure 2**). B cells had a diverse group of outgoing and incoming signals and may possibly downregulate outgoing ESAM pathways and incoming SELL pathways (**Figure 2a and Figure 2b**). Endothelial cluster and CAF clusters had increased outgoing ESAM and PERIOSTIN signaling (**Figure 2a and Figure 2b**).

**Figure 2.**
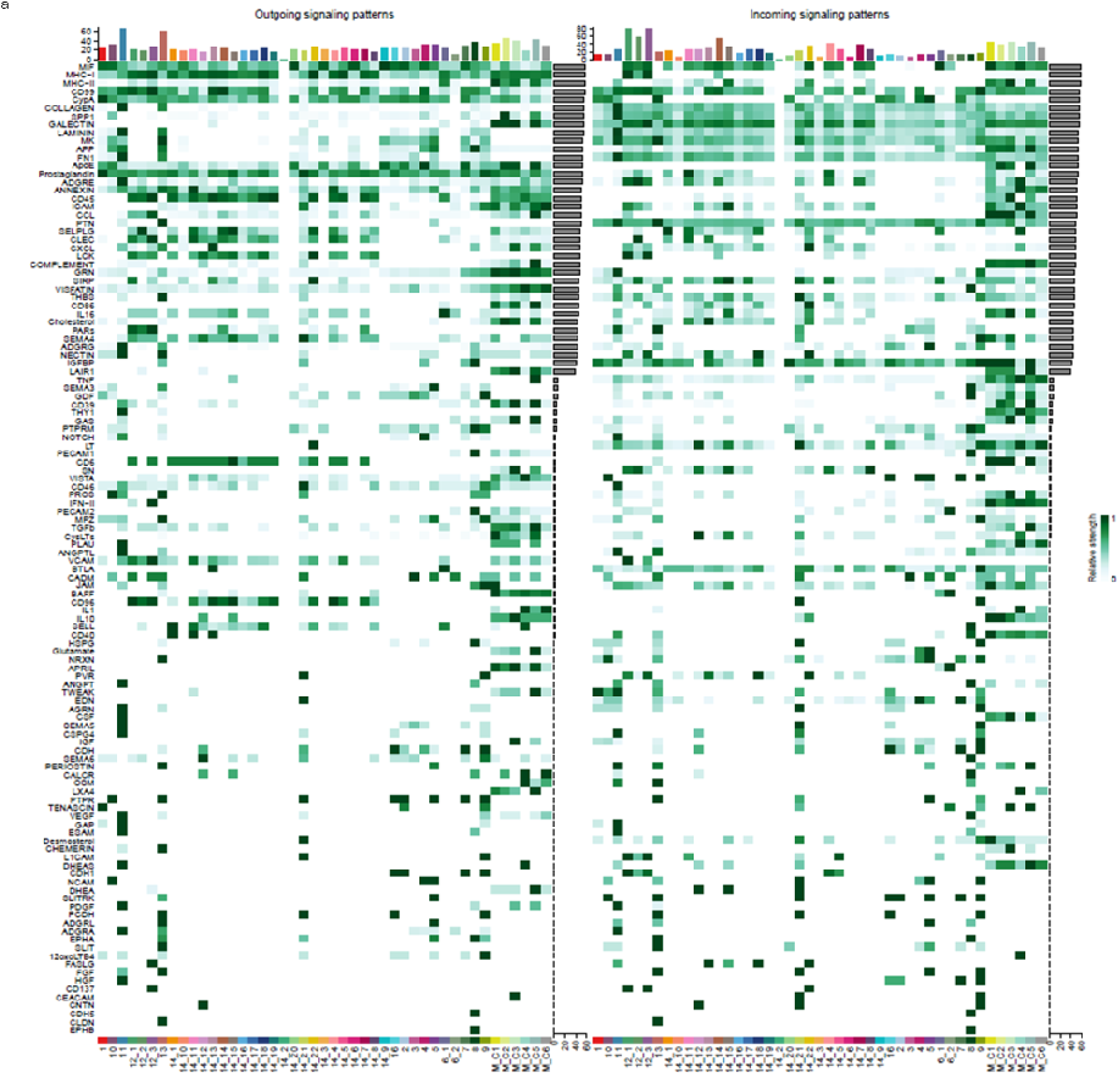
Overall Signal Motif by Cell Chat Analysis of each Cell Type Cluster. **a,** Outgoing signal pattern and incoming signal pattern depicted by CellChat Analysis with depiction of signal motif by cell type cluster. T cells showed increased MHC-I outgoing signaling (inflammatory signaling) and myeloid cells upregulated LAIR1 and APRIL (inflammatory signaling).

### Myeloid Immune Cell Associated with Survival in Metastatic Melanoma

Kaplan-Meier curves were generated, p=0.029, (**Figure 3a**) based on the results and cut-off values determined by percentage infiltration of total sample on recursive partitioning analysis (RPA) (**Figure 3b**). The only significant difference between groups with treatment factors was an increased rate of targeted therapy in the best and worse survival groups (31% vs 5.3% vs 43%, p=0.036) in **Figure 3c**. In addition, although 22 of 42 patients had response treatment with available data the best survival group still had 56% of patients with No Response (NR) in **Figure 3c**. On cox regression analysis Node 5 vs Node 2 was significantly different, HR 4.24, 95% CI 1.4-12.9, p=0.011 in **Figure 3d**. However, Node 4 vs Node 2 approached significance, HR 1.89, 95% CI 0.82-4.25, p=0.13 in **Figure 3d**. Median Overall Survival (OS) months differed between Node 2 at 52 vs Node 4 at 32 vs Node 5 as 13 in **Figure 3e**.

**Figure 3.**
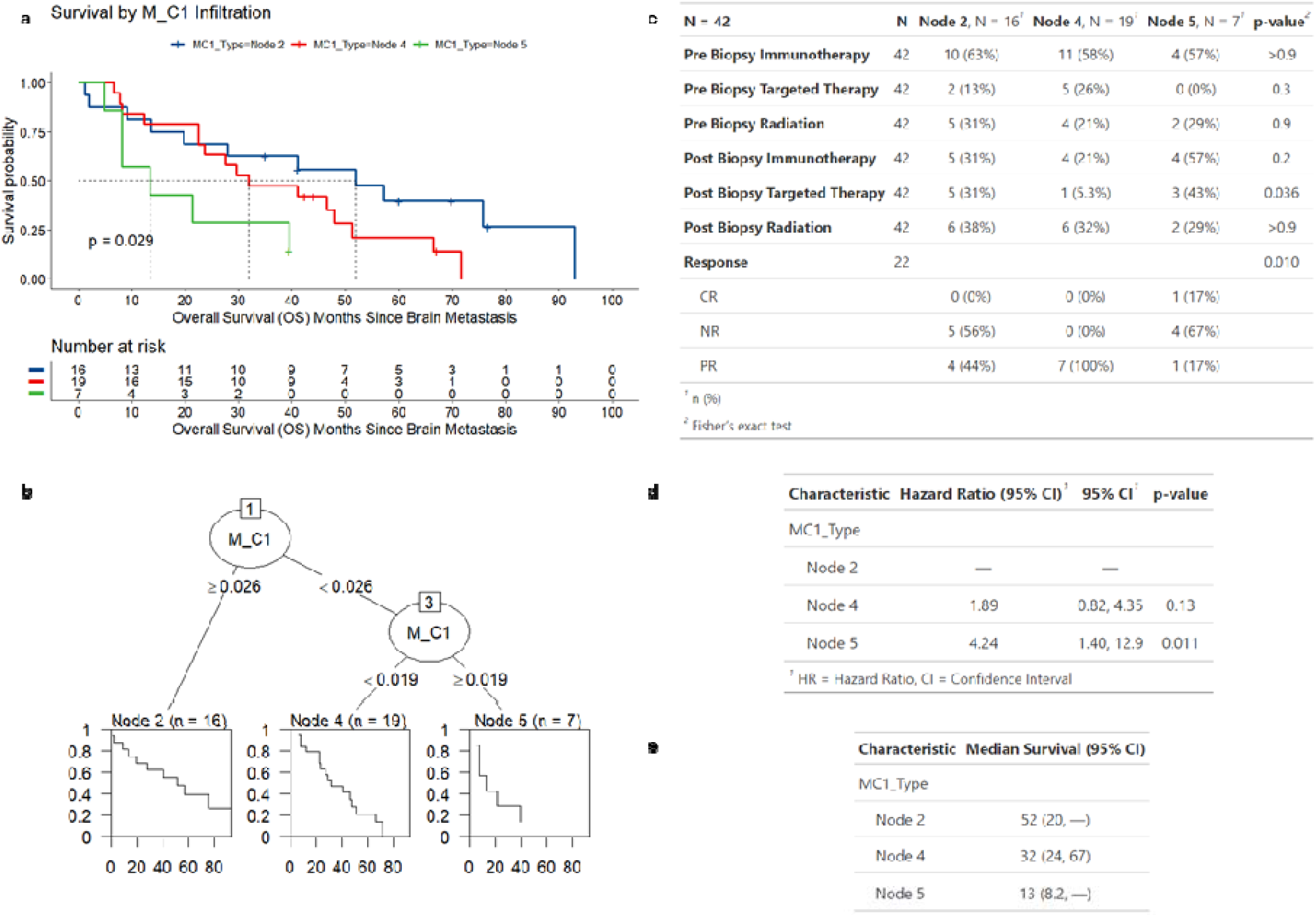
Myeloid Cell Type 1 (M_C1) Infiltration Improves Overall Survival (OS) in Metastatic Melanoma. **a,** Kaplan-Meier survival curve depicting overall survival (OS) for brain metastases cohort when stratified by M_C1 cell frequency identified by recursive partitioning analysis (RPA). **b,** RPA results of training brain metastases cohort depicting each “Node” identified with cut-off frequency of M_C1 cell determined by analysis. **c,** Table depicting patient characteristics between RPA groups with univariate analysis conducted. **d,** Cox regression analysis of RPA groups on OS with calculated Hazard Ratio. **e,** Median OS months calculated for each RPA group.

### Interaction of Myeloid Immune Cell and Other Cell Types in Metastatic Melanoma

Across statistical tests the following cell types were significantly different between groups: M_C1, M_C3, M_C4, 9, and M_C6 (**Supplement 2**). Overall, it appears that myeloid cell types appear to be the most significant prognostic factor between each group. In addition, Cluster 9 appears to be highest in Node 5, with the worst OS outcomes and increased expression of AC015802.6, PRMT5, TROVE2, PYCR1, and MINOS1 compared to other clusters. Other than tumor cells, the worst OS group (Node 5) has lower M_C1 and M_C3 clusters but similar monocytes and regulatory macrophage type cells (**Supplement 2**).

### External Validation of Myeloid Cell Type on Survival in Advanced Melanoma & Cell Interaction

After calculating Kullback-Leibler (KL) Divergence for each model against the distribution of cell types in our original scRNA sequencing dataset, the CIBER methodology produced results that most closely matched the distribution expected of cell types on the data from the nivolumab clinical trial data set (**Supplement 3**).The nivolumab clinical trial data set included a cohort of advanced melanoma patients with pre-treatment biopsy and on-treatment biopsy bulk RNA sequencing while receiving nivolumab. Pre-treatment biopsy M_C1 infiltration scores did not correlate with OS (**Figure 4a & Figure 4b**) p=0.71 while on-treatment biopsy scores predicted OS (**Figure 4c & Figure 4d**) p=0.048. A shift in M_C1 infiltration scores was clinically but not statistically significant while undergoing treatment with nivolumab in **Figure 4e**, p=0.0695.

**Figure 4.**
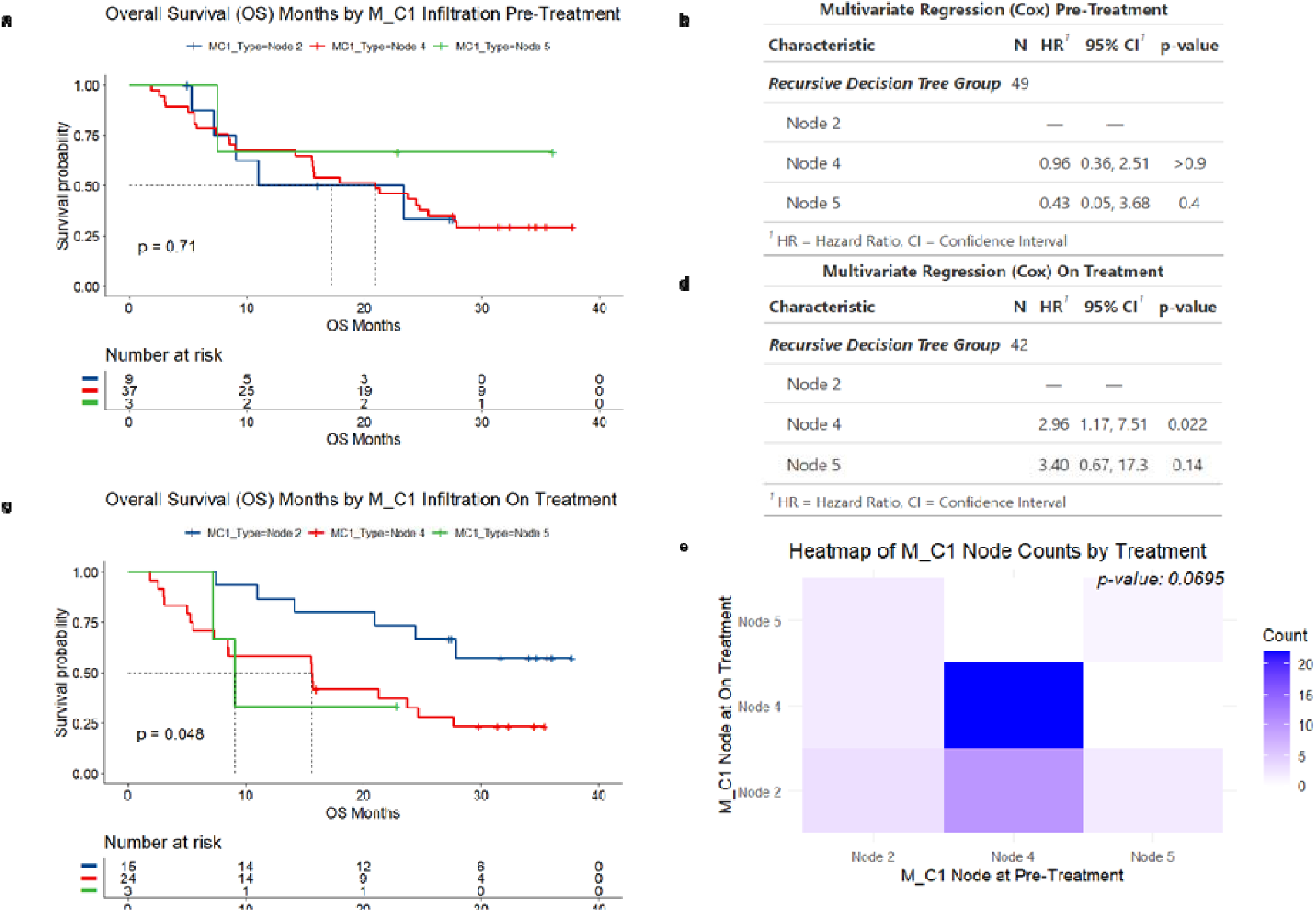
M_C1 Infiltration is Affected by Nivolumab Treatment in Pre vs On Treatment Biopsy and Improved Overall Survival (OS) **a,** Kaplan-Meier plot of Overall Survival (OS) Months for patients stratified by M_C1 infiltration nodes based on pre-treatment biopsy samples with bulk RNA sequencing with cell type percentages calculated by deconvolution using CIBERSORT. **b,** multivariate cox regression analysis of M_C1 infiltration nodes for pre-treatment biopsy sequencing **c,** OS by stratification of M_C1 infiltration nodes for on treatment biopsy samples **d,** multivariate cox regression analysis of M_C1 infiltration nodes for on treatment biopsy sequencing **e,** Stuart-Marxwell test of treatment effect of Nivolumab on pre-treatment to on treatment node assignment based on bulk RNA sequencing results analyzed by deconvolution using CIBERSORT

Differences in frequencies in individual cell types in each pre-treatment, on-treatment, and paired treatment sample were analyzed with statistical testing. For pre-treatment biopsies, M_C1 groups were associated with significant differences in clusters 14_19 and M_C1 across statistical testing (**Supplement 4**). Without accounting for response on nivolumab, high M_C1 infiltration scores were associated with low 14_19 (early CD3) (**Supplement 4**).

For on-treatment biopsies M_C1 groups were associated with significant differences in clusters 12_1, 14_2, 14_6, 2, 6_2, 9 & M_C1 across statistical testing (**Supplement 5**). Tumor clusters 2 and 9 negatively correlated with OS and 14_2 positively correlated with OS. High 12_1 (NKT Activated with PD1), lower 14_6 (early CD3), and higher 6_2 (B Cell Type 2) were associated with improved M_C1 infiltration and OS. This points to both a cellular and humoral response in the immune system in predicting improved OS based on M_C1 infiltration.

Furthermore, paired patient samples for pre-treatment biopsies were analyzed by changes from M_C1 infiltration groups by three categories while undergoing treatment: Better, Stable, and Worse. When evaluating pre-treatment biopsies, patients have poor response to nivolumab and subsequent M_C1 infiltration when starting with high M_C1 and 12_3 (PD1+NKT) cells (**Supplement 6**). However, when 14_9 (early CD3) was high, patients improved on nivolumab (**Supplement 6**). Analysis of on-treatment biopsies was inconclusive.

### External Validation of Myeloid Cell Type on Survival in TCGA Melanoma & Cell Interaction

After calculating Kullback-Leibler (KL) Divergence for each model against the distribution of cell types in our original scRNA sequencing dataset, the CIBER methodology produced results that most closely matched the distribution expected of cell types on the TCGA data set (**Supplement 3**).

Kaplan-Meier plots were calculated by stage of melanoma at diagnosis (**Figure 5a**), by M_C1 infiltration for all patients (**Figure 5b**), M_C1 infiltration for Stage I (**Figure 5c**), M_C1 infiltration for Stage II (**Figure 5d**), M_C1 infiltration for Stage III (**Figure 5e**), and M_C1 infiltration for Stage IV (**Figure 5f**) all with significant p-values. Taken together, it appears that M_C1 infiltration groups are prognostic of OS in the TCGA. There appears to be a paradoxical relationship of Node 5 patients with worse OS in Stage I and complex interactions with OS in Stage II/III/IV. However, the highest and lowest infiltration e.g. Node 2 and Node 4 appear to be prognostic on all Stages of melanoma (**Figure 5**).

**Figure 5.**
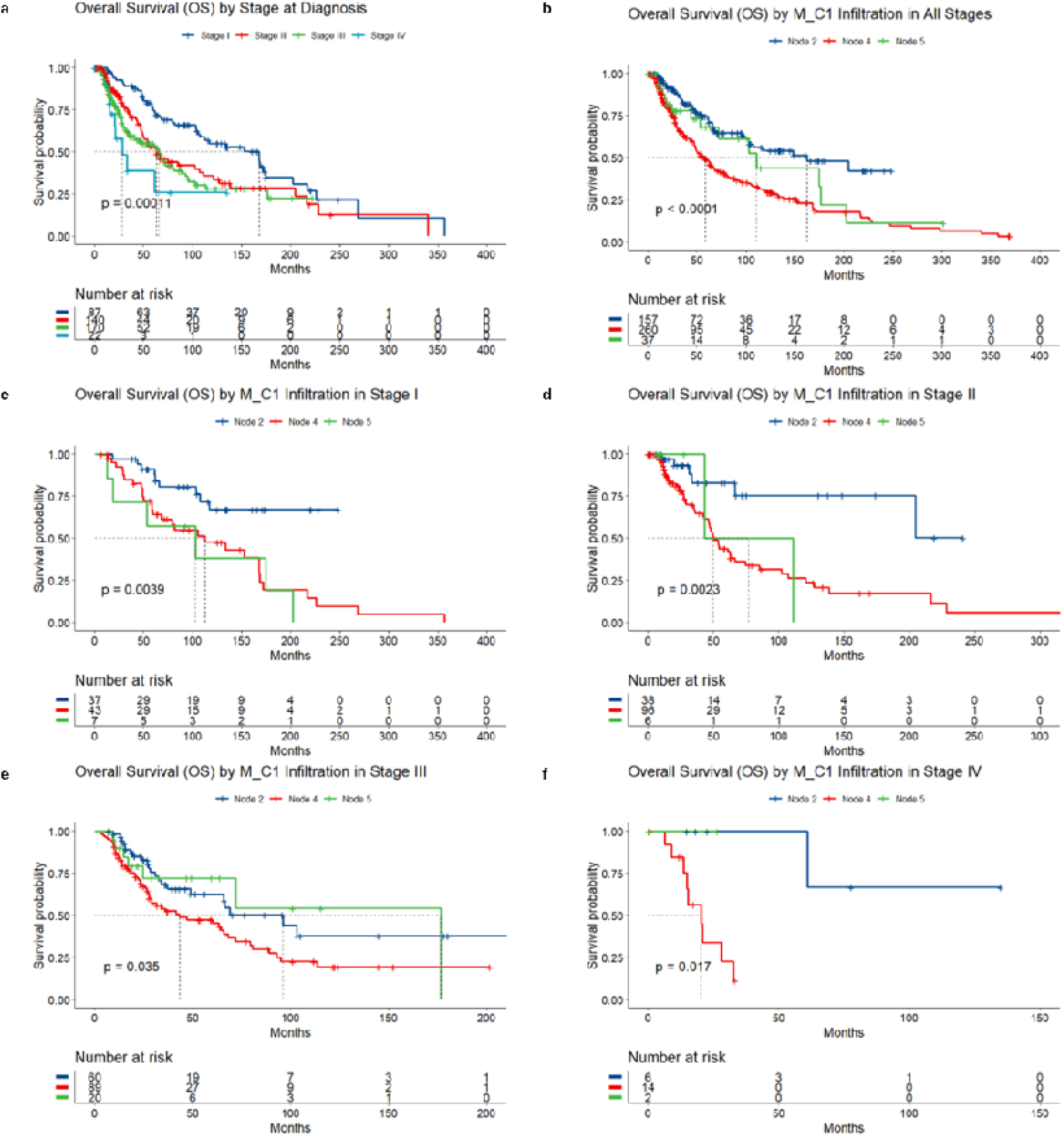
Overall Survival (OS) in TCGA Cohort by Stage of Melanoma and M_C1 Infiltration. **a-f,** Kaplan-Meier analysis of Overall Survival (OS) months stratified by stage of melanoma at diagnosis (a) and M_C1 infiltration nodes by Stage of melanoma (b-f).

Analyzing differences between infiltration groups across statistical testing found 21 cell types that had significance across the three statistical methods (**Supplement 7**). Only cluster 9 tumor cells showed significant differences between all three infiltration groups in **Supplement 7**. 12_1 (NKT) was significantly different between all three groups in **Supplement 7**. Clusters 14_4 (CD8 T Cell Exhausted) and 14_7 (CD8 T Cell Exhausted with PD1) were both significantly increased in Node 4 vs the two other groups in **Supplement 7**. Cluster 14_6 (Naïve CD4 1) was highest in Node 4 and 14_17 (Naïve CD4 3) highest in Node 2 with differences between all groups in **Supplement 7**. Cluster 14_22 (CD4 Treg 2) was highest in Node 2 with no difference between Node 4 and 5 in **Supplement 7**.

Overall, this points to high M_C1 infiltration being associated with less malignant tumor (Cluster 9), increased cytotoxic NKT vs exhausted T cell (12_1 vs 14_4/14_7), increased T reg (14_22), and B Cell infiltration (6_1/6_2). This supports an immune mediated hypothesis directed by increased antigen presentation by a DC-macrophage (M_C1) cell leading to both cellular and humoral immune response as shown by increased cytotoxic NKT Cells (12_1) and B Cells (6_1/6_2).

## Discussion

A total of 45 cell types including tumor cells (13), fibroblasts (2), endothelial cells (1), NK cells (3), CD3 T cells (4), CD4 T cells (11), CD8 T cells (3), B cells (2), and myeloid cells (6) were found after clustering with scCCESS. When analyzing differences between major cell groups by tissue type (e.g. primary melanoma, acral, brain metastases, etc.) only myeloid cells showed no differences in frequency by tissue type **Figure 1**. Overall, tumor cells showed heterogeneity within tissue types as well as some tissue specific elevations in tumor clusters which supports both tissue specificity in melanoma sub-types and consistency for melanoma in **Supplement 1**. Immune cell subsets between tissue types varied; however, immune cell subsets between primary melanoma vs non-brain metastatic melanoma differed in the following clusters: 14_6 (early CD4), 14_8 (early CD4), 14_11 (CD4 + GATA3), 14_16 (early CD3), 14_19 (early CD3), and 6_1 (B Cell Type 1) in **Supplement 1**. Generally, this reflects more systemic immune activation in metastatic melanoma compared to primary melanoma. CellChat analysis appeared consistent with understanding of immune cluster function, although tumor cell clusters were heterogenous in signaling pathways.

When examining the myeloid cell clusters on OS in the brain metastases subset of patients, M_C1 was the only cell type with significant clinical association based on the infiltration cut-off scores found using RPA. Interestingly, high infiltration (Node 2) had the best survival; medium infiltration (Node 5) had the worst survival, and low infiltration (Node 4) appeared to be in the middle survival curve. Other than M_C1 infiltration only M_C3 (Macrophage Inflammatory Type 2) appeared to be significantly different between all three groups in **Supplement 2**. Other between group differences appear to be driven by tumor heterogeneity (Cluster 2, 9, and 10), M_C4 (monocyte), M_C6 (Macrophage Regulatory), and 14_16 (early CD3) in **Supplement 2**.

The model was externally validated in a clinical trial data set using bulk RNA sequencing with a deconvolution method with minimized KL Divergence using our scRNA labeling as a map for cell identification. When analyzing a subset of patients treated with nivolumab with matched biopsies prior to treatment and while on treatment, our model showed significant differences by M_C1 infiltration node when assessing the on-treatment biopsy in **Figure 4**. No other cell types other than M_C1 were significantly different between all three groups in **Supplement 5**. However, between group differences with agreement in all three analyses included tumor heterogeneity (Cluster 2, 9, and 14_2), 12_1 (NKT Cell Type 1), 14_6 (early CD3), and 6_2 (B Cell Type 2) in **Supplement 5**. This shows consistency in tumor cell type when compared to the brain metastases data, but includes other novel mechanisms associated with NKT and B Cells. This correlation supports the importance of checkpoint activity and therapeutic targeting across myeloid populations, with a possible mechanism of M_C1 function as a DC-Macrophage to increase B cell response. When evaluating changes in response groups, the treatment effect of nivolumab was not significant in **Figure 4**, p=0.069. 4 of 7 patients (57%) with high M_C1 infiltration on pre-treatment biopsy had a shift to lower infiltration on nivolumab. 12 of 35 patients (34%) shifted from a lower infiltration group into high infiltration Node 2 while 23 patients did not change M_C1 infiltration (66%).

When externally validating the model using the TCGA, our results showed the M_C1 was prognostic across all stages in the high (Node 2) and low (Node 4) groups in **Figure 5**.

However, Node 5 appears to be prognostic as the worst OS only in Stage I melanoma and does not appear to be significantly different when compared to Node 2 in other stages of melanoma. Given the ability to predict OS in Stage I melanoma despite including this in the model, M_C1 infiltration provides a novel method of evaluating early stage melanoma.

Our identified cell type, M_C1, appears to be a novel hybrid cell exhibiting markers of macrophages and dendritic cells. Biological development shows these cells are related with terminal differentiation in each sub-type^40^. Interestingly, new subtypes of DCs have been discovered in both blood and tissue in humans with a recent finding Villani et al^41^. Furthermore, hybrid cells such as Natural Killer – T cells (NKT) have been previously described and it may also be the case that hybrid DC-Macrophages may also be identifiable^42,43^. This may be explained by the idea of lineage immune cell plasticity which is supported by a recent cell trajectory analysis by Wei et al^44^. Overall, this finding underscores the restriction of underclassifying macrophage phenotypes and behaviors using single or binary nomeclatures. Limited, positive-selection based labeling strategies may have failed to identify hybrid-type cells of importance and have likely contributed to both the current lack of clarity in TAM behaviors and historically limited descriptions of significant TAM differences in checkpoint-refractory disease.

Reanalyzing public scRNA seq data for melanoma has provided additional information due to larger sample sizes and alternative analysis techniques which include: BRAF mutations^31^, NK cytotoxicity^32^, a four gene ferroptosis related prognostic signature^33^, CD8 T cell function^34,35^, monocyte derived prognostic signature^36^, NKT and CD4 T cell related survival improvements^37^, and CAF associated metastasis and response to immune therapy^38^. Our finding of a novel cell type (M_C1) with prognostication of OS across all stages of melanoma provides the possibility of unlocking biomarkers and cell therapies in the future. In addition, our clustering methodology should be viewed as highlighting significant gene expression across a normal distribution. Some cells within in the cluster may be more macrophage like whilst others may resemble DCs more on the extremes of gene expression within the cluster. Validation using flow cytometry and immunohistochemical staining of human melanoma using genes or markers with high expression in our model may provide further evidence of this cell type within the tumor microenvironment in melanoma and its role in tumor resistance. In addition, isolating and evaluating cellular function of the M_C1 cluster could prove to provide a novel cell therapy in the future that unlocks a convergence of innate and adaptive immune response against tumors.

## Methods

### Data extraction

We collected five single-cell RNA sequencing (scRNA-seq) datasets and two bulk RNA sequencing datasets for humans from publicly available repositories, including Gene Expression Omnibus (GEO)^14,18,29,30,45^, Single Cell Portal^17^, and The Cancer Genome Atlas (TCGA)^46^. For the 10X Genomics data, raw sequencing files were obtained using SRA-Toolkit (v3.0.2) via the fastq-dump utility. After obtaining the raw sequencing data, gene expression (GEX) matrices were generated through a comprehensive pipeline. First, the data were demultiplexed to separate individual cell barcodes from sequencing reads. Next, barcode processing was performed to filter out low-quality or ambiguous barcodes, ensuring accurate assignment of reads to specific cells. The processed reads were then aligned to the human reference genome Hg38 using a highly optimized alignment algorithm. Finally, gene quantification was carried out, whereby aligned reads were counted to generate a matrix of gene expression levels across all cells. All steps in this pipeline were performed using the 10X Genomics CellRanger software (v7.1.0). For datasets generated using the SMART-Seq2 protocol, GEX matrices were directly downloaded from the Gene Expression Omnibus (GEO) and Single Cell Portal repositories.

### Single-cell RNA Data Filtering and Normalization

To enhance the accuracy of gene expression estimates, we utilized the ‘remove-background’ function in CellBender (version 0.3.0) for each sample.^47^ CellBender utilizes a deep learning-based approach to model and remove technical artifacts, such as background noise and ambient RNA, which can contaminate single-cell RNA sequencing data. By estimating the true biological signal from the raw gene-by-cell count data, CellBender generates ambient-corrected count matrices that provide a more accurate reflection of the cellular transcriptome.

These corrected matrices were then imported into R (version 4.2.2) and converted into Seurat objects using Seurat (version 4.9.9.9086).^48^ We applied stringent cell selection criteria to ensure data quality: cells were retained if they exhibited less than 10% or up to 20% mitochondrial reads, expressed between 3,500 and 7,500 genes and had unique molecular identifier (UMI) counts ranging from 500 to 60,000. Next, doublet cells were identified using Scrublet (version 0.2.3)^49^, which computes doublet scores based on the expected doublet rate, preset at 10%. Cells flagged as doublets by Scrublet with the default parameters were excluded from further analysis.

For datasets generated using the SMART-Seq2 protocol, we incorporated pre-filtered datasets sourced from public repositories, including GEO and Single Cell Portal.

After data filtering, the gene-by-cell expression matrices were merged using the ‘mergè function. We then proceeded with pre-processing using SCTransform (version 0.4.0)^50^, which incorporates a variance-stabilizing transformation (VST) method. SCTransform replaces conventional normalization and scaling procedures with a regularized negative binomial regression model, effectively controlling for technical noise. This method adjusts gene expression measurements by modeling unwanted variation and emphasizing biological variability.

### Unsupervised dimensional reduction and clustering

For dimensionality reduction, Principal Component Analysis (PCA) was conducted. PCA reduces the dimensionality of the data by transforming the original variables into a new set of variables, which are linear combinations of the original variables. These new variables, called Principal Components (PCs), are ordered so that the first few retain most of the variation present in the original variables.

Upon completion of PCA, the datasets were prepared for integration using Harmony^51^, applied through the ‘RunHarmony’ function with specified covariates: project, sequencing techniques, tissues, and donors. Harmony leverages a model-based approach to adjust the principal components across different datasets, mitigating the impact of batch effects and other technical discrepancies. This step ensures that the integrated data are aligned in a shared dimensional space that more accurately reflects the underlying biological heterogeneity.

Following the successful integration of datasets, we employed Uniform Manifold Approximation and Projection (UMAP) for further dimension reduction. We used the top 30 corrected principal components derived from Harmony. UMAP operates on a manifold learning technique that maps high-dimensional data into a more manageable two-dimensional space, facilitating the visualization and interpretation of complex data structures and relationships.

### Clustering and identification of marker genes

To determine the optimal number of clusters across the entire cell population, we utilized the ‘estimate_k’ function from scCCESS (version 0.3.3)^52^, configuring the criteria method to use NMI criterion and selecting the k-means algorithm. We set the ensemble size to 10 to ensure robustness by averaging results across multiple initializations. Once the optimal cluster number was identified, we applied it in the ‘ensemble_cluster’ function, again using the k-means algorithm, to partition the cells into distinct clusters based on their gene expression profiles.

Subsequently, we focused on the myeloid cell population. The clustering process was repeated specifically for these cells. We began with the ‘estimate_k’ function, using the same parameters as those applied to the whole cell population. We then utilized ‘ensemble_cluster’ with the k-means algorithm and the newly determined optimal number of clusters to better capture the unique expression patterns of the myeloid cells. This independent analysis resulted in a different optimal number of clusters, reflecting the distinct biological characteristics of the myeloid cells. This procedure was repeated on three additional clusters representing B cells, NK cells, and T cells.

Next, we identified significant marker genes for each cell subset using the ‘FindMarkers’ function in Seurat. We employed the MAST^53^ method for statistical testing, which is well-suited for single-cell RNA-seq data, particularly in handling sparsity and zero-inflation. To ensure robustness in the marker identification process, we set the minimum percentage (min.pct) of cell expressing a gene to 10%, meaning that a gene had to be expressed in at least 10% of cells in either of the groups being compared. Additionally, we applied the logarithmic fold change threshold (logfc.threshold) of 0.1, focusing on genes that exhibited at least a modest difference in expression levels. These criteria allowed us to identify marker genes that are both statistically significant and biologically relevant for each subset of cells.

### CellChat Analysis

We used CellChat version 2.1.2^54^ for cell-cell communication computation to assess signaling changes using our defined cell types

### Deconvolution of Bulk RNA Datasets for Cell Type Estimation

Integrated single cell dataset with our cell type labels was used to deconvolute bulk RNA-seq data of TCGA cutaneous melanoma. Estimation of cell proportion were computed using Please add the three methodologies here. I will also had we optimized model choice by KL Divergence for the final comparison. CIBERSORTx version 1.0.1^55^, BayesPrism version 2.2.2^56^, and MuSiC version 1.0^57^. Models were compared for fit based on Kullback-Leibler Divergence in R version 4.3.2 for all 3 deconvolution methods.

### Recursive Partitioning Analysis

Recursive partitioning analysis (RPA) was conducted in R version 4.3.2 with package rpart. Myeloid cell types M_C1 to M_C6 were evaluated with prognostic implications on overall survival (OS) on a cohort of n=42 samples with complete data with melanoma brain metastases. Duplicated samples (e.g. more than 1 biopsy at 1 timepoint for a patient) were averaged in percent expression for analysis.

### Survival Analysis

Survival analysis was conducted in R version 4.3.2 using packages survival and survminer. Values of M_C1 infiltration from RPA analysis were used as thresholds for risk groups termed ‘nodes’ for analyses on the following cohorts: Melanoma brain metastases, nivolumab clinical trial data set, tumor cancer genome atlas (TCGA) melanoma. Kaplan-Meier curves were generated and statistical testing using log-rank analysis and cox regression analysis was used to compared differences between groups.

### Statistical Analysis Between Groups

Groups were tested for differences based on clinical factors using Fisher’s exact test. For treatment effect of nivolumab on risk ‘node’ the Stuart-Maxwell test was conducted. For differences in frequencies of cell types based on cell type clusters in the training and validation data cohorts the following statistical tests were conducted: ANOVA, Kruskal-Wallis with Benjamini-Hochberg correction, and Wilcoxon Rank Sum with Benjamini-Hochberg correction. All analyses were conducted in R version 4.3.2.

### Reporting Summary

## Data availability

For the single cell RNA sequencing datasets used in this analysis, the raw FASTQ files can be downloaded from the Gene Expression Omnibus (GEO), https://www.ncbi.nlm.nih.gov/geo/, under the accession number GSE189889, GSE215120 and GSE174401, and gene expression matrices are available from GEO under accession number GSE115978 and GSE185386, and from Single cell Portal, https://singlecell.broadinstitute.org/single_cell, under SCP number SCP1493. Seurat object for the Integrated dataset including gene expression matrices, UMAP coordinates, cell type labels, and other cell metadata is available at Figshare: 10.6084/m9.figshare.28196423. For the TCGA bulk RNA sequencing, the raw gene expression matrix can be downloaded from GEO under the accession number GSE62944.

## Code availability

All code used was publicly available.

## Acknowledgements

We would like to thank the hard work and dedication of all the groups that we acquired publicly available data from in order to conduct our analysis.

## Author contributions

MUA, LH, RN, and AK contributed equally to the conception, design, analysis, and development of the manuscript for publication.

SS and EM provided support on the analysis and development of the manuscript.

## Additional information

Supplementary information is included as Supplement 1 through Supplement 8 as supporting information for this work.

Correspondence should be addressed to Amanda Kirane at.

