## Supplementary figures and images for "Single Cell RNA Sequencing Analysis of Human Melanoma Reveals A Distinct Prognostic Myeloid Cell"

### Supplement 1

#### Slide 1
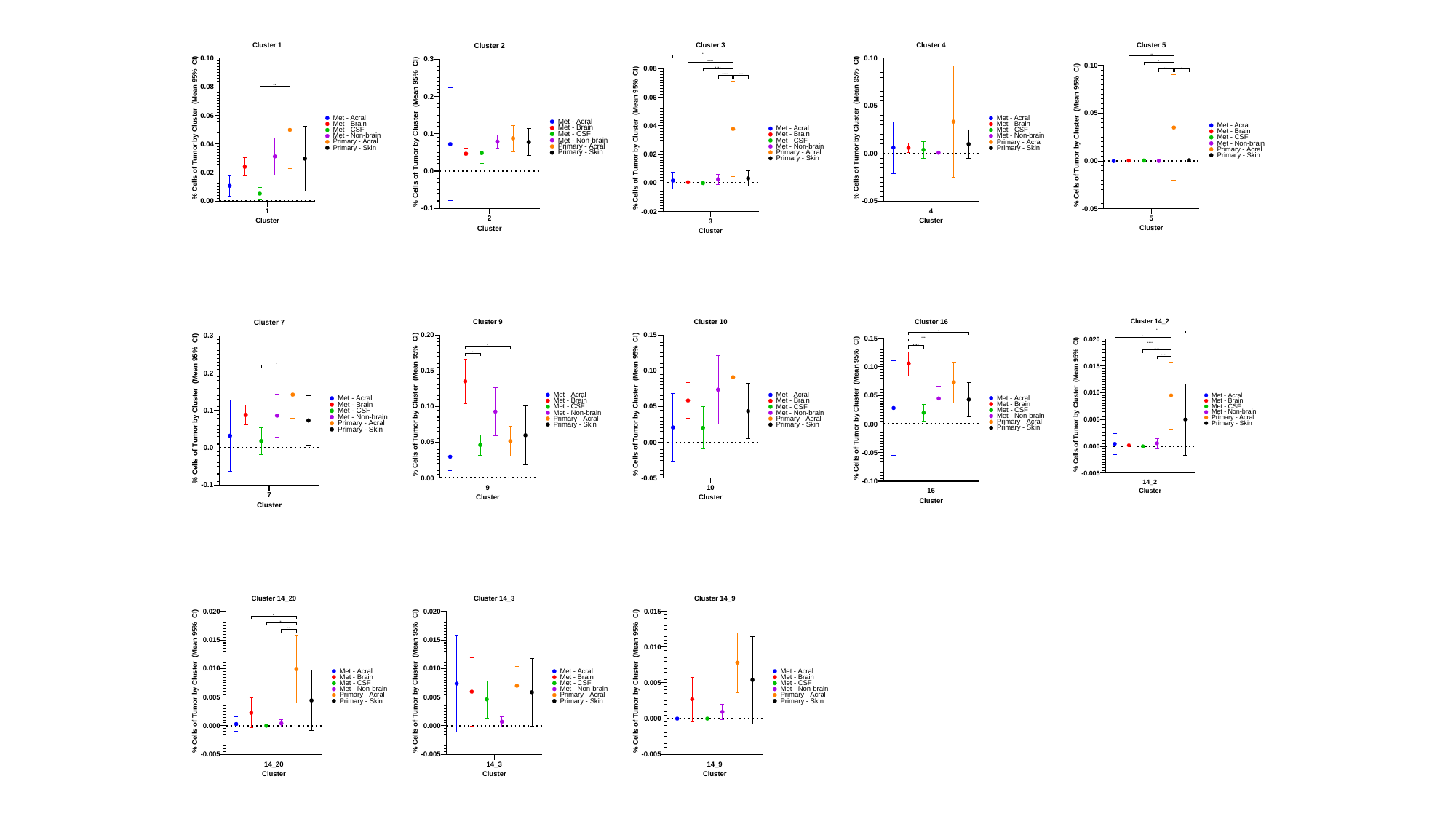

#### Slide 2
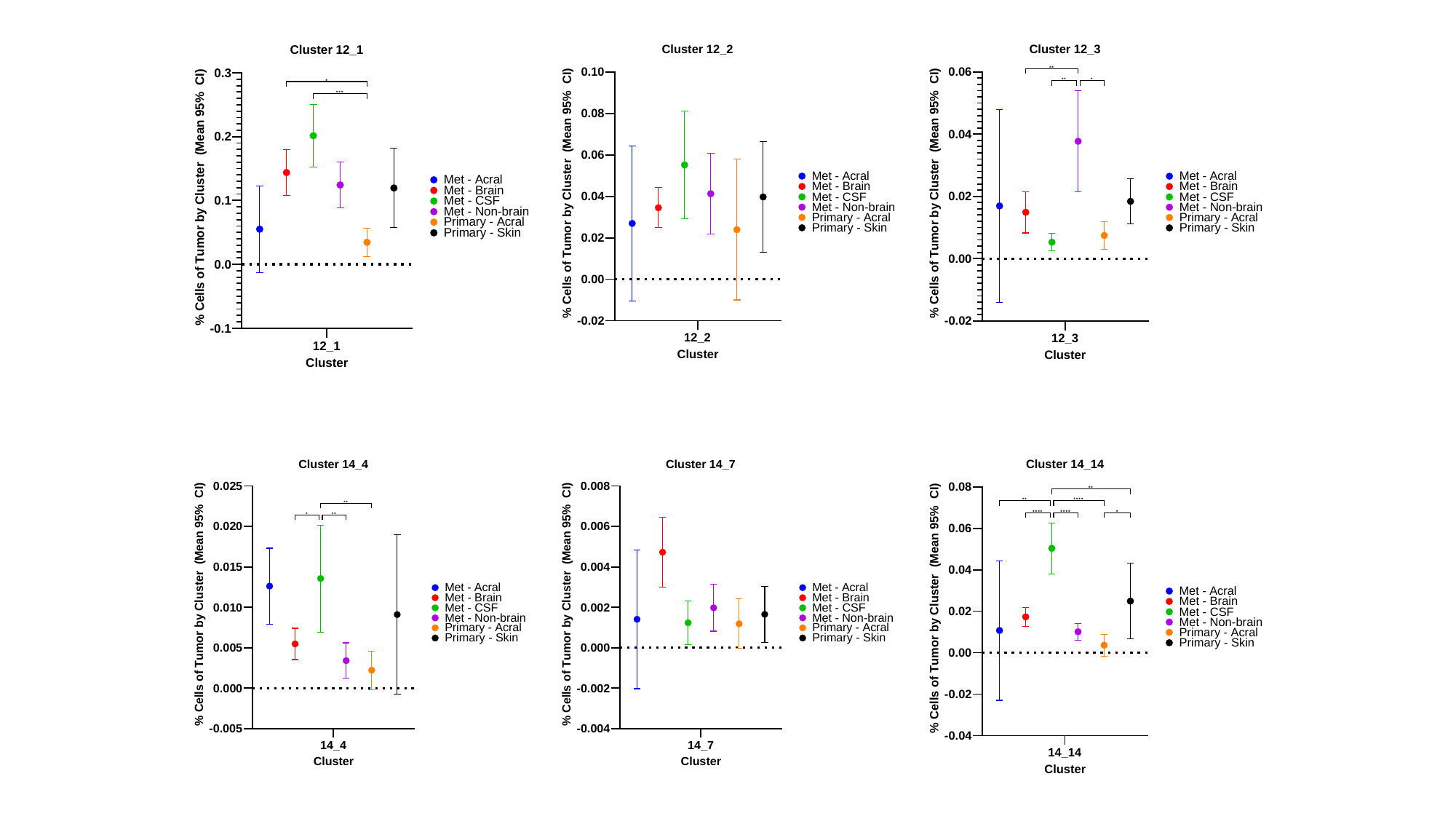

#### Slide 3
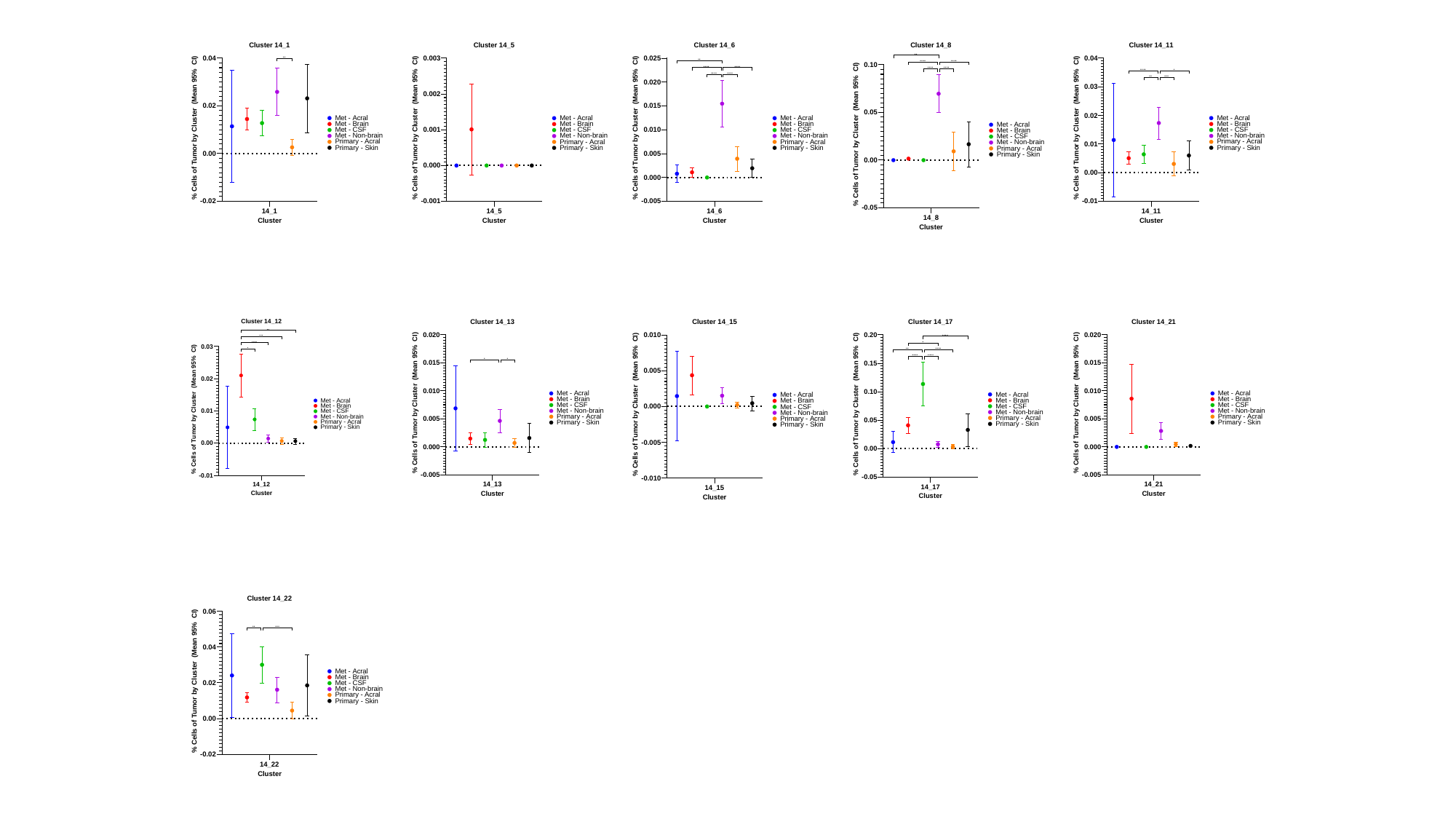

#### Slide 4
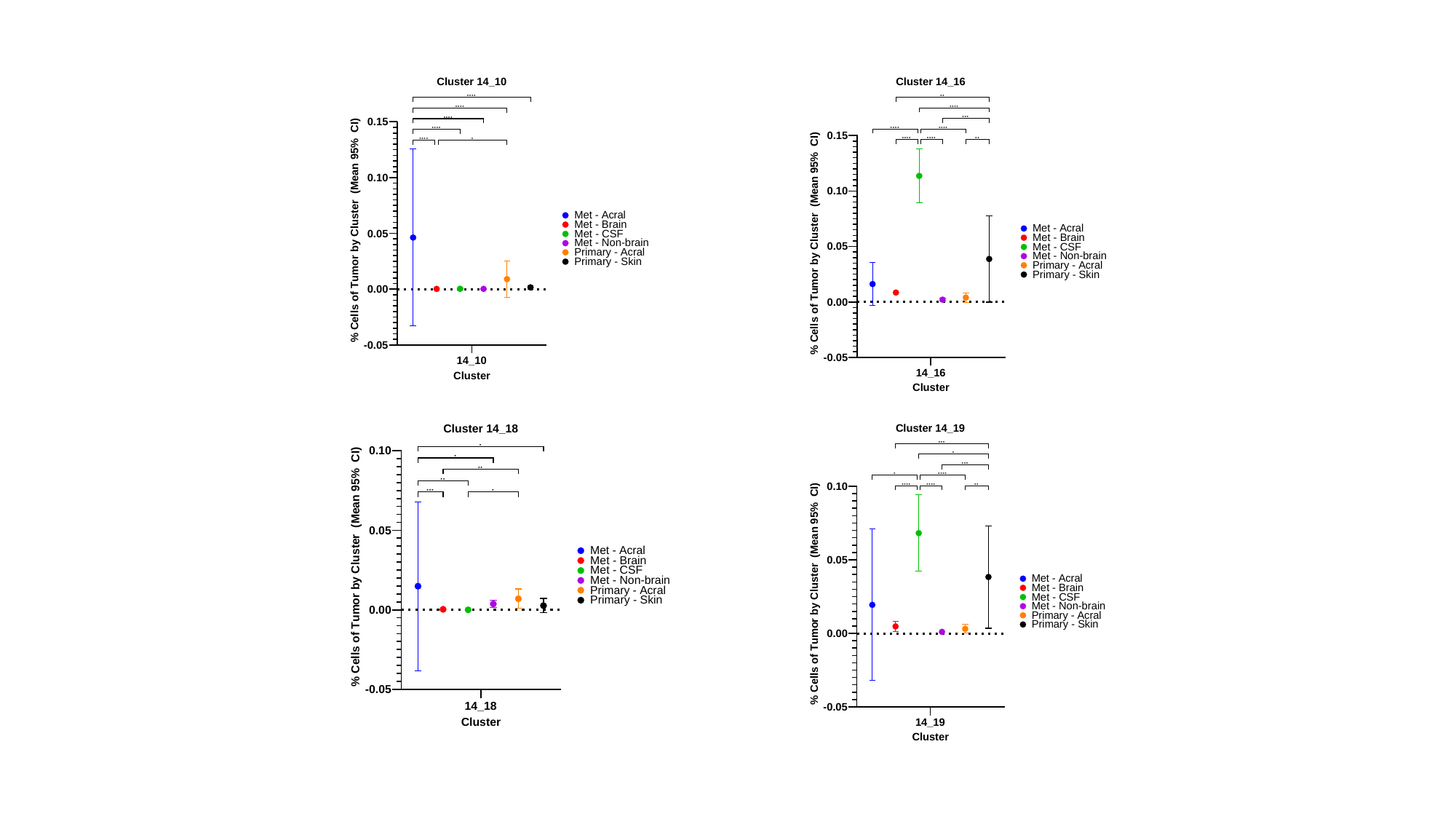

#### Slide 5
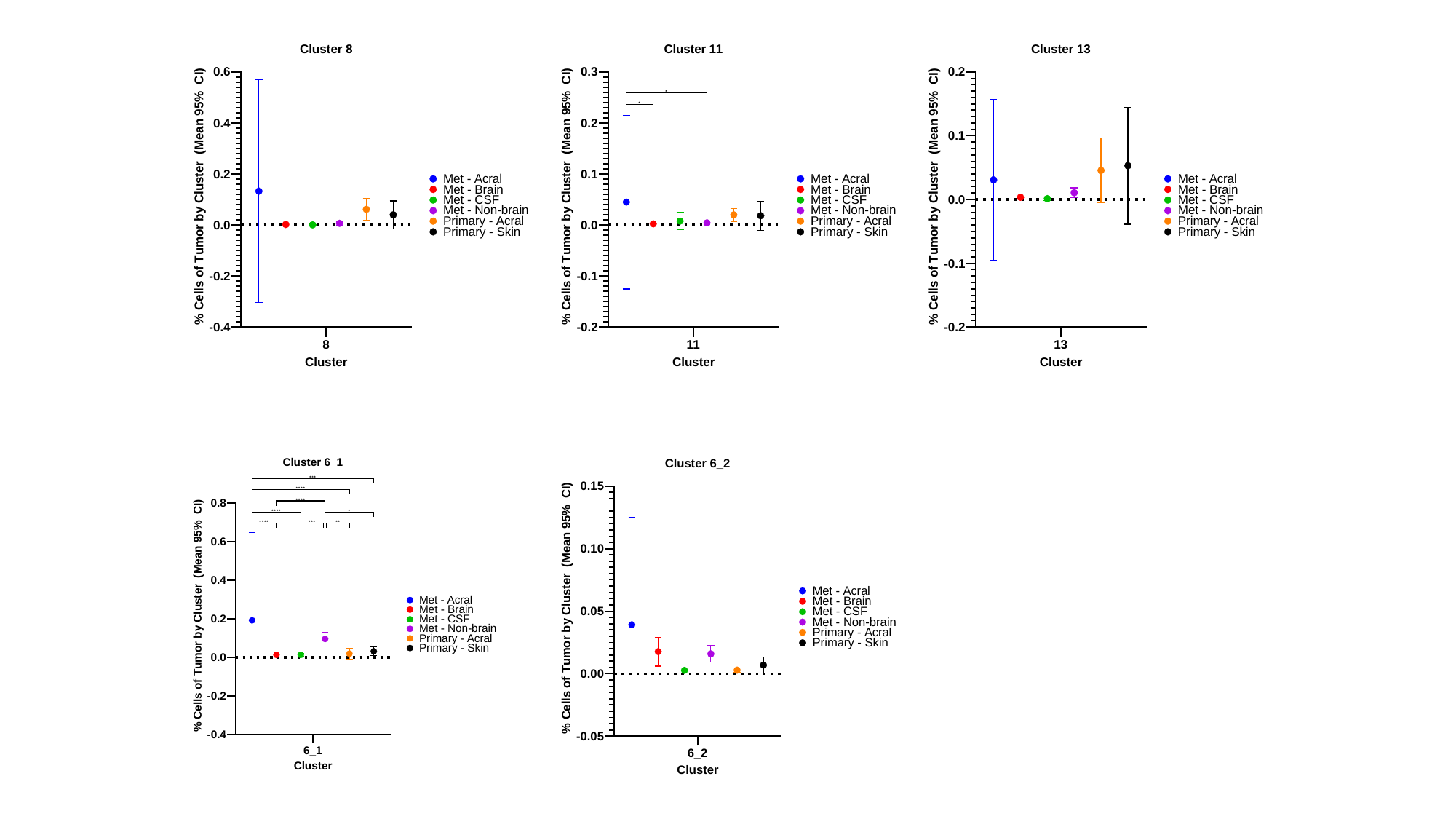

#### Slide 6
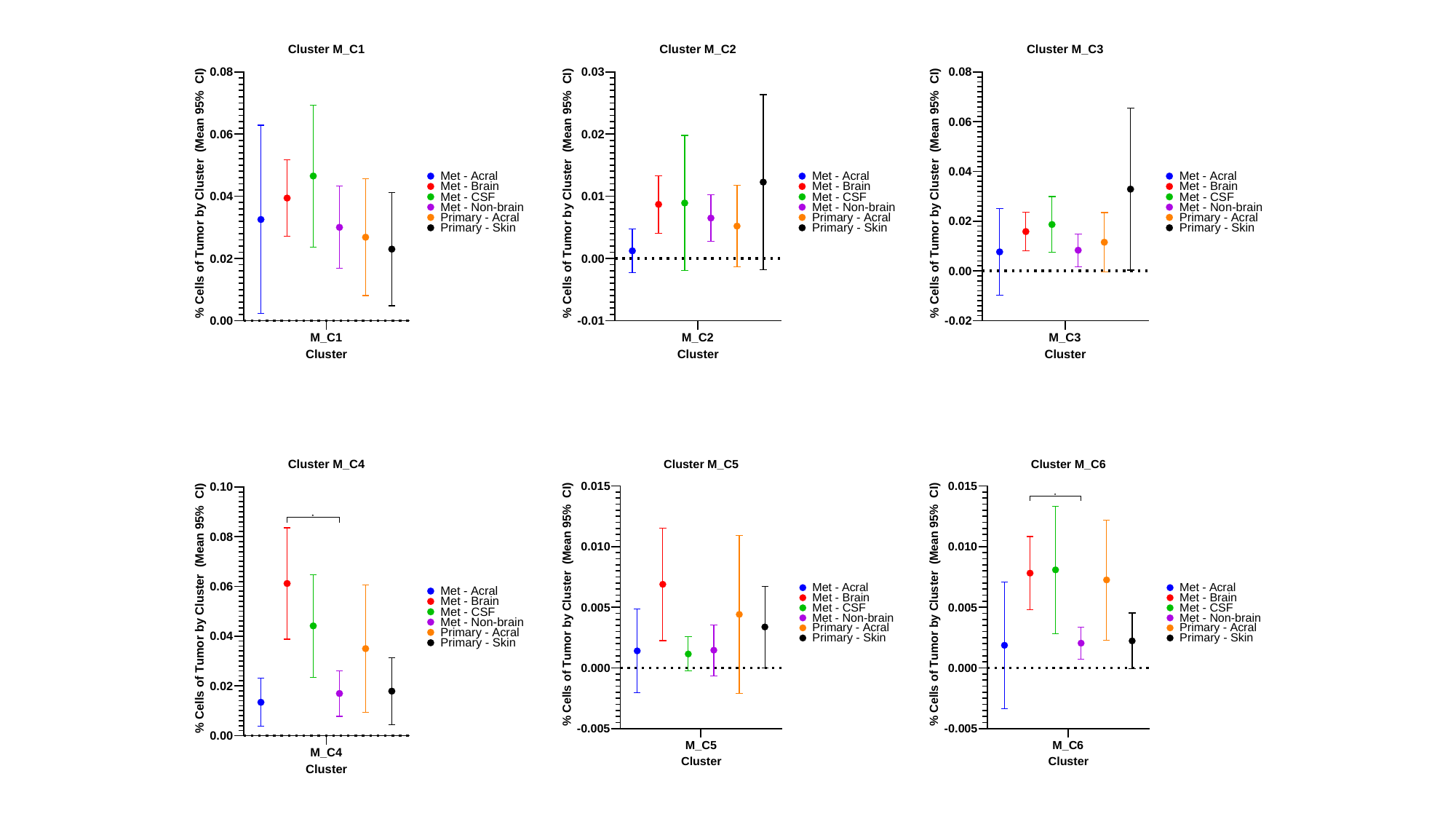

### Supplement 2

#### Slide 1
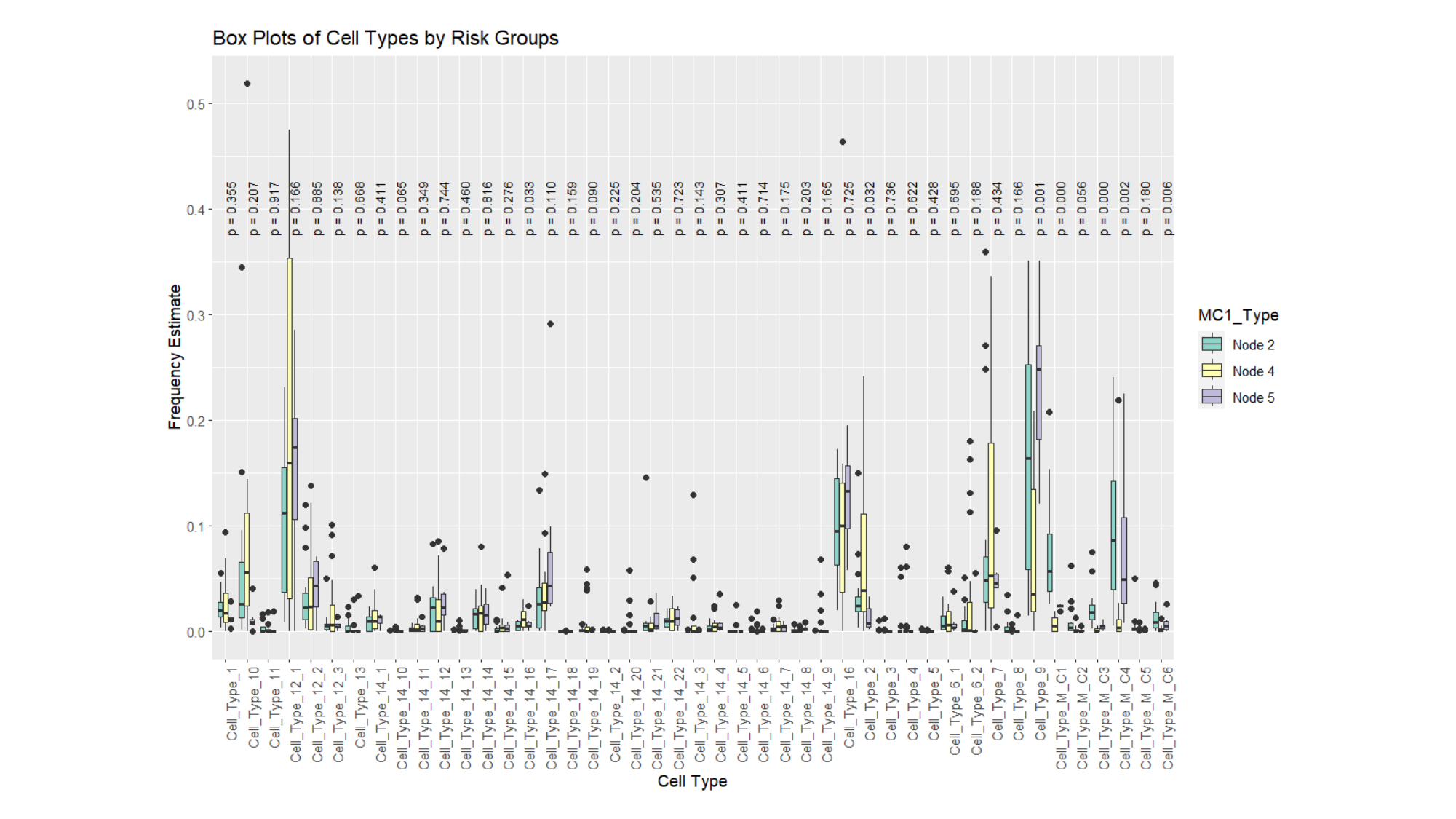

#### Slide 2
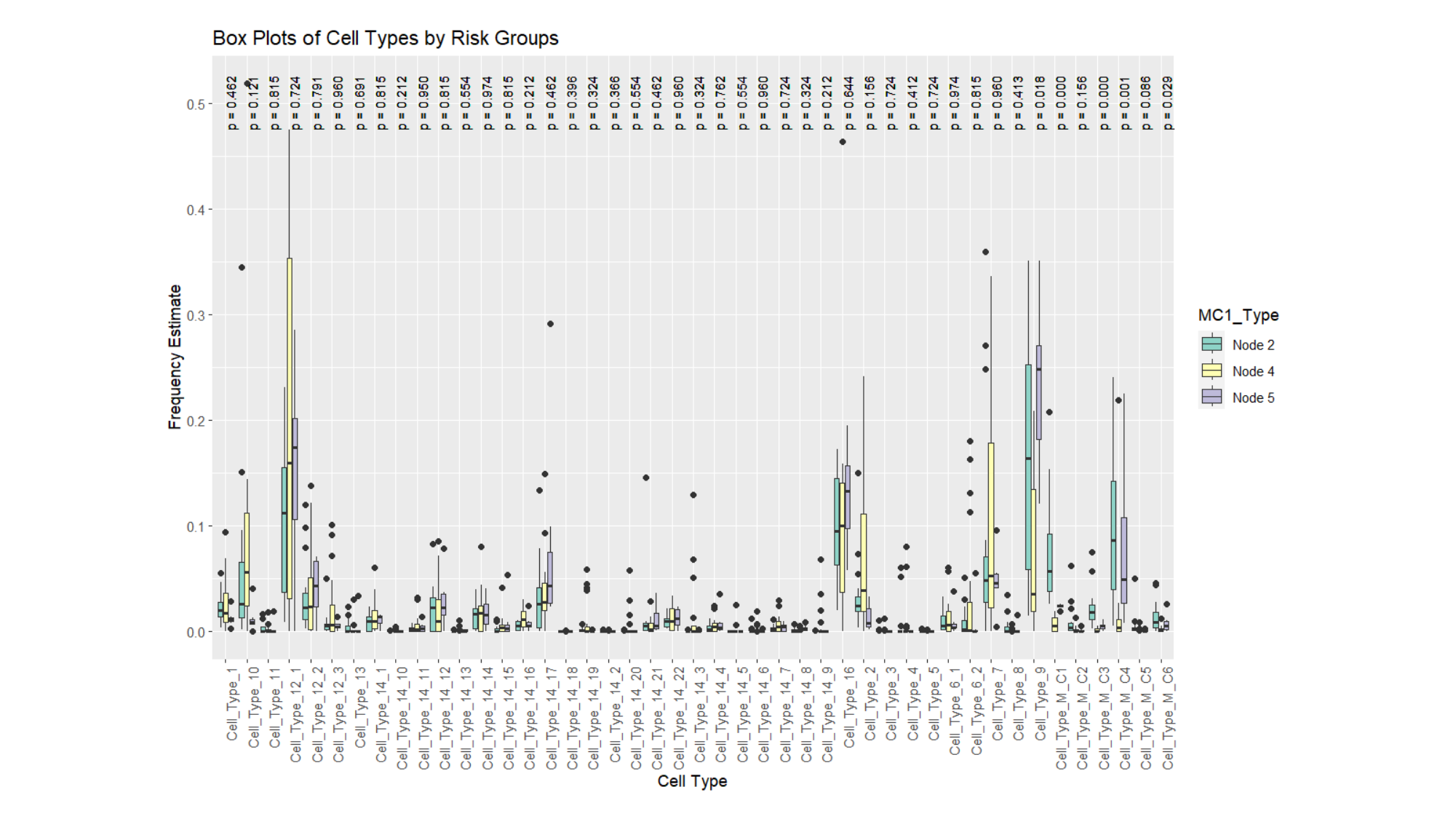

#### Slide 3
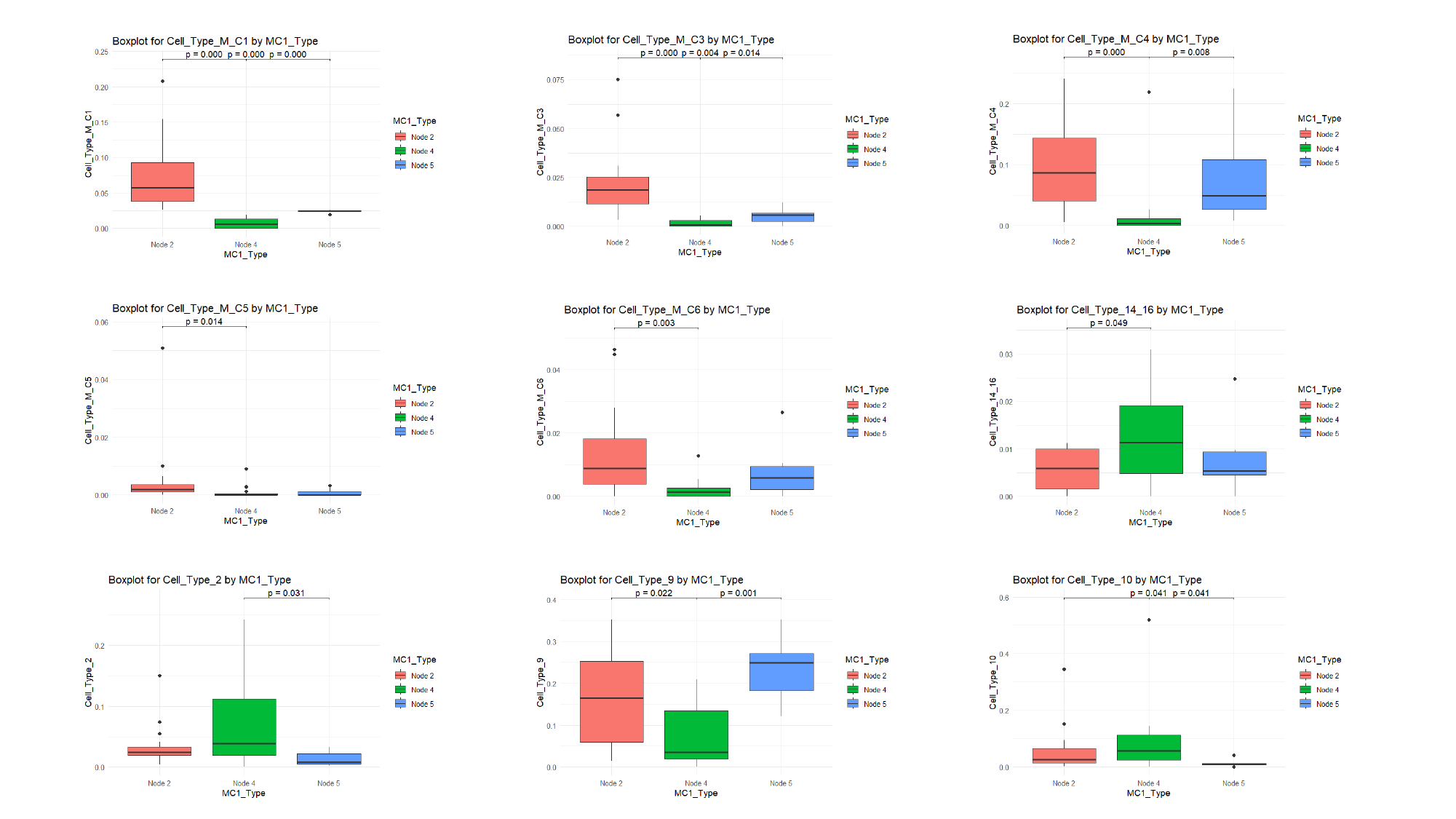

### Supplement 4

#### Slide 1
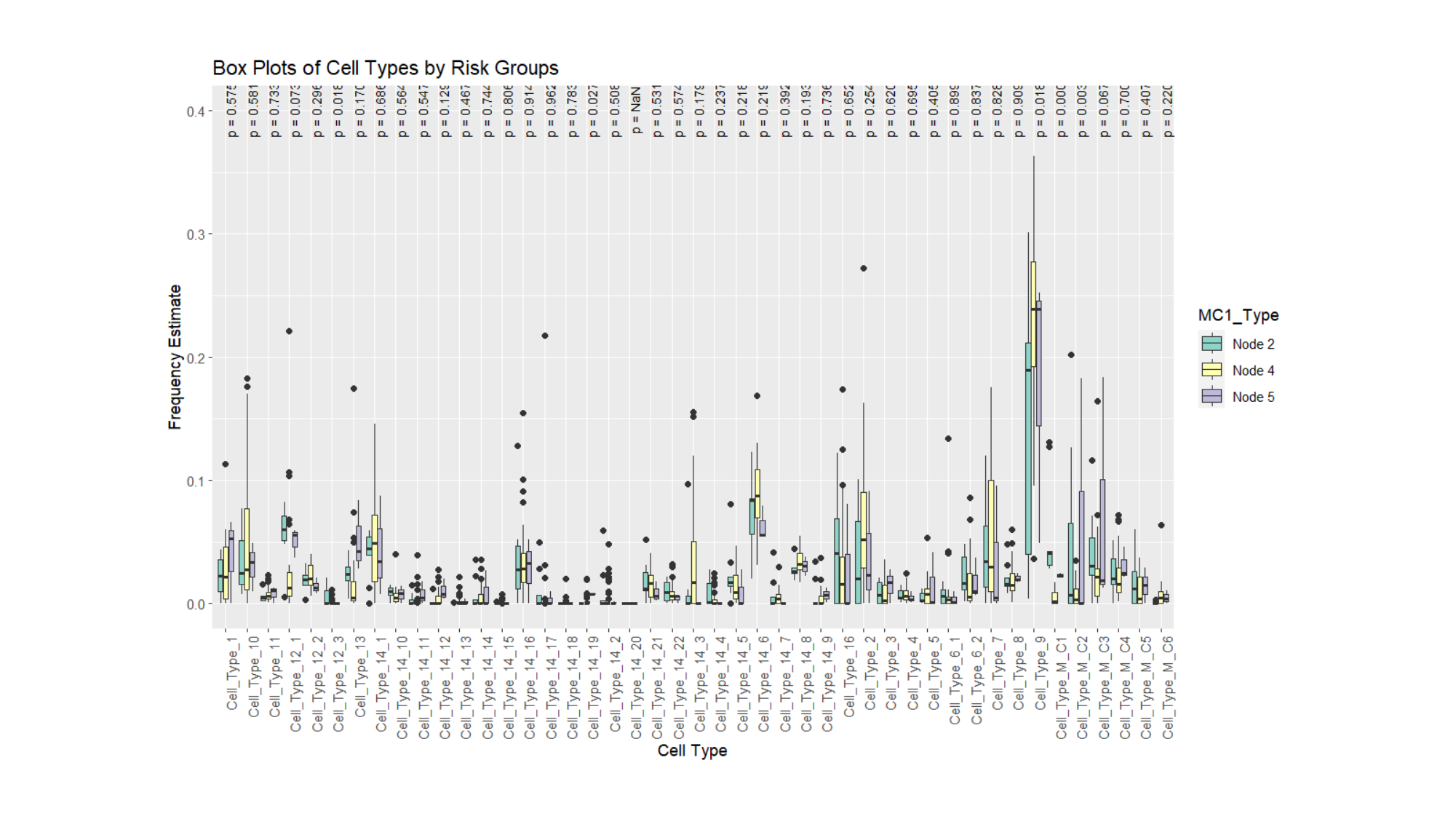

#### Slide 2
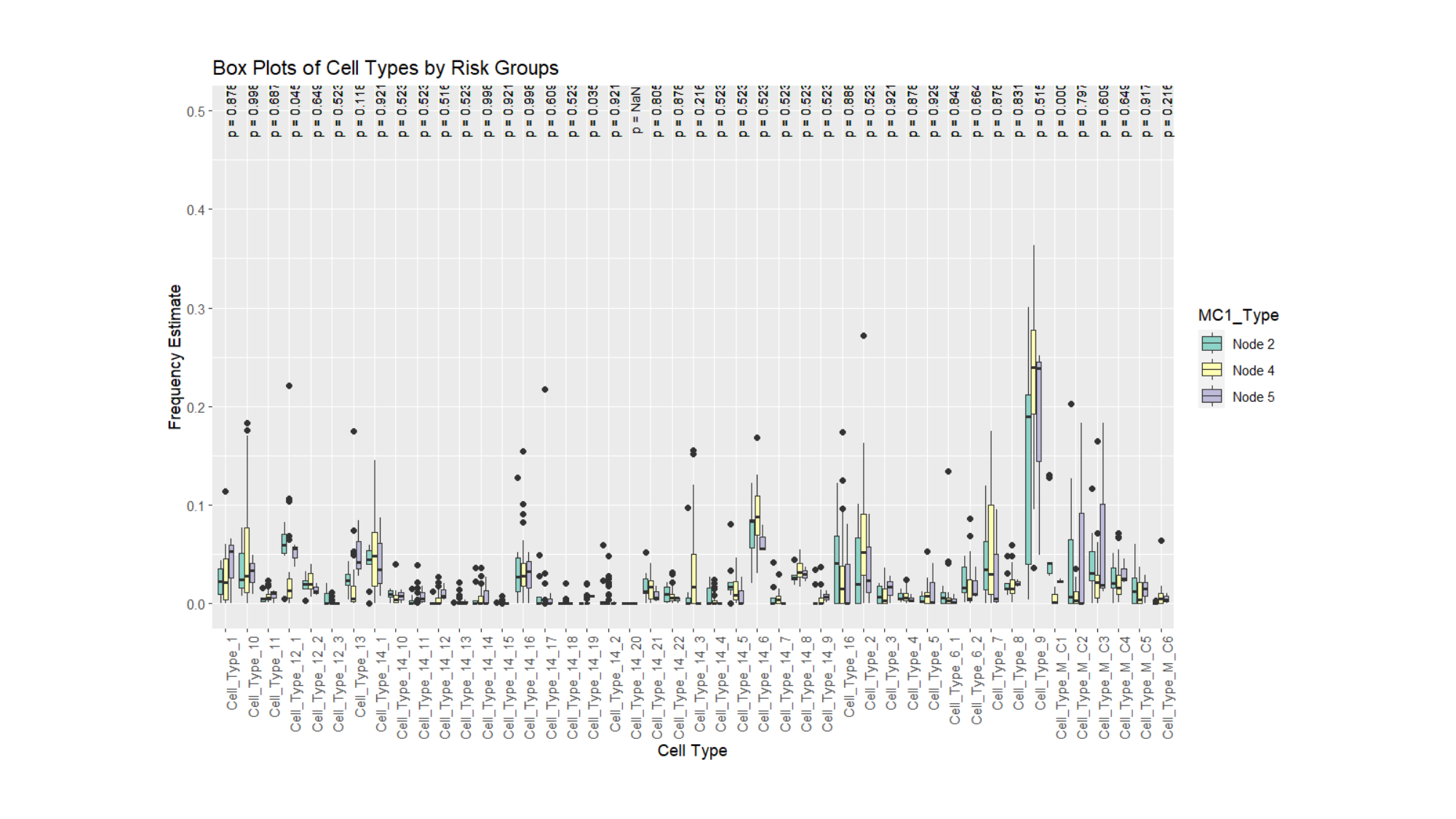

#### Slide 3
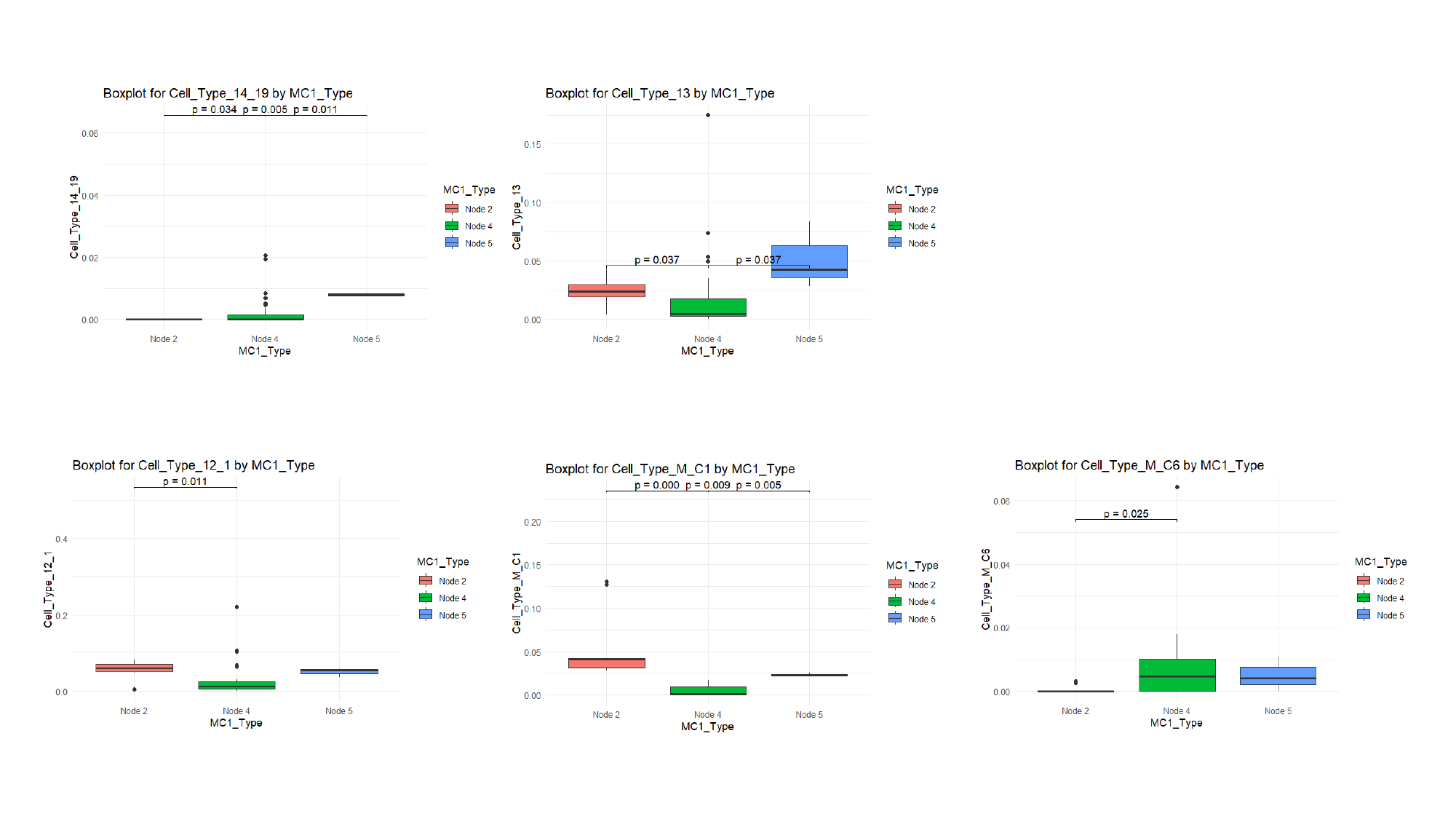

### Supplement 5

#### Slide 1
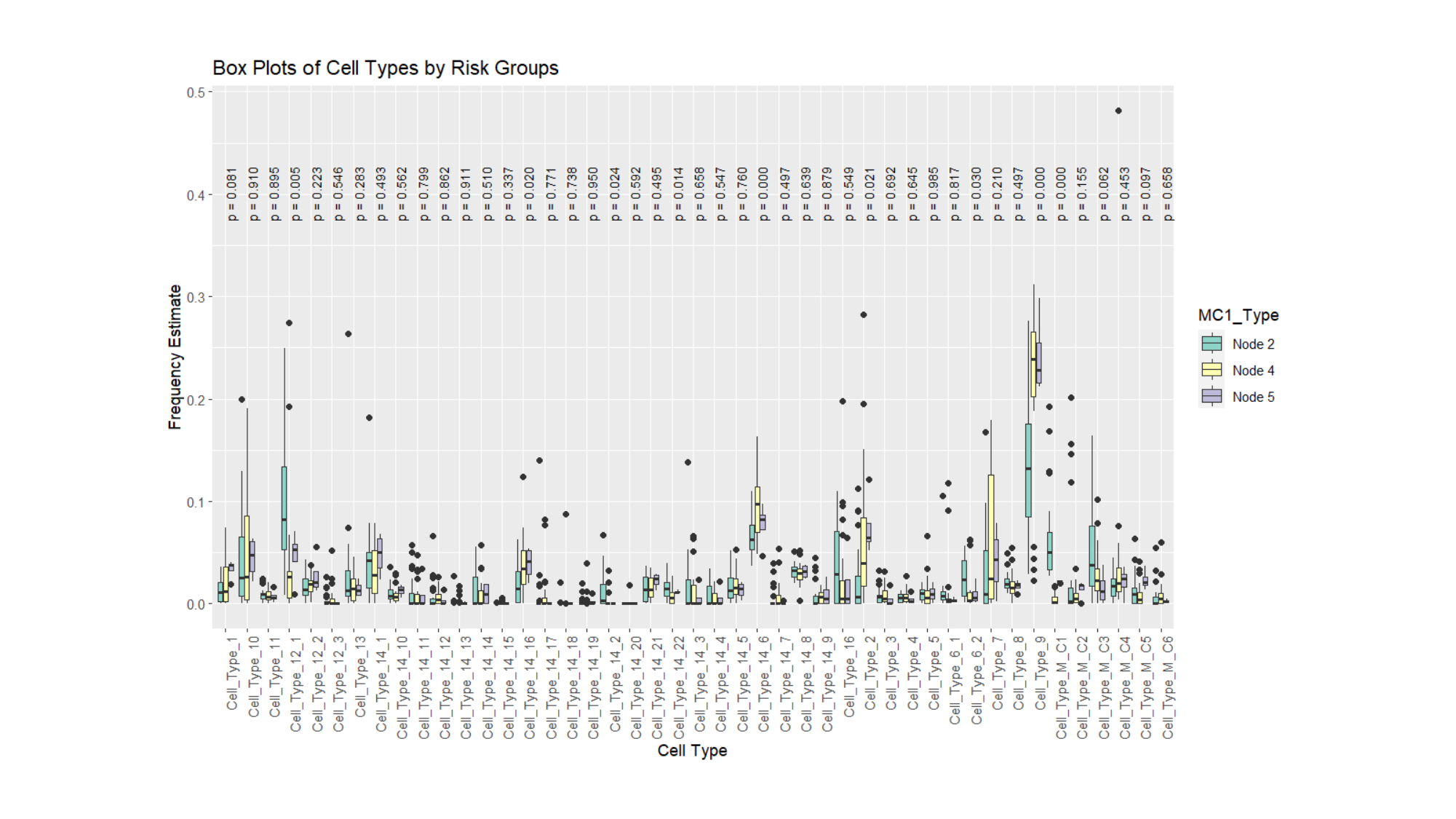

#### Slide 2
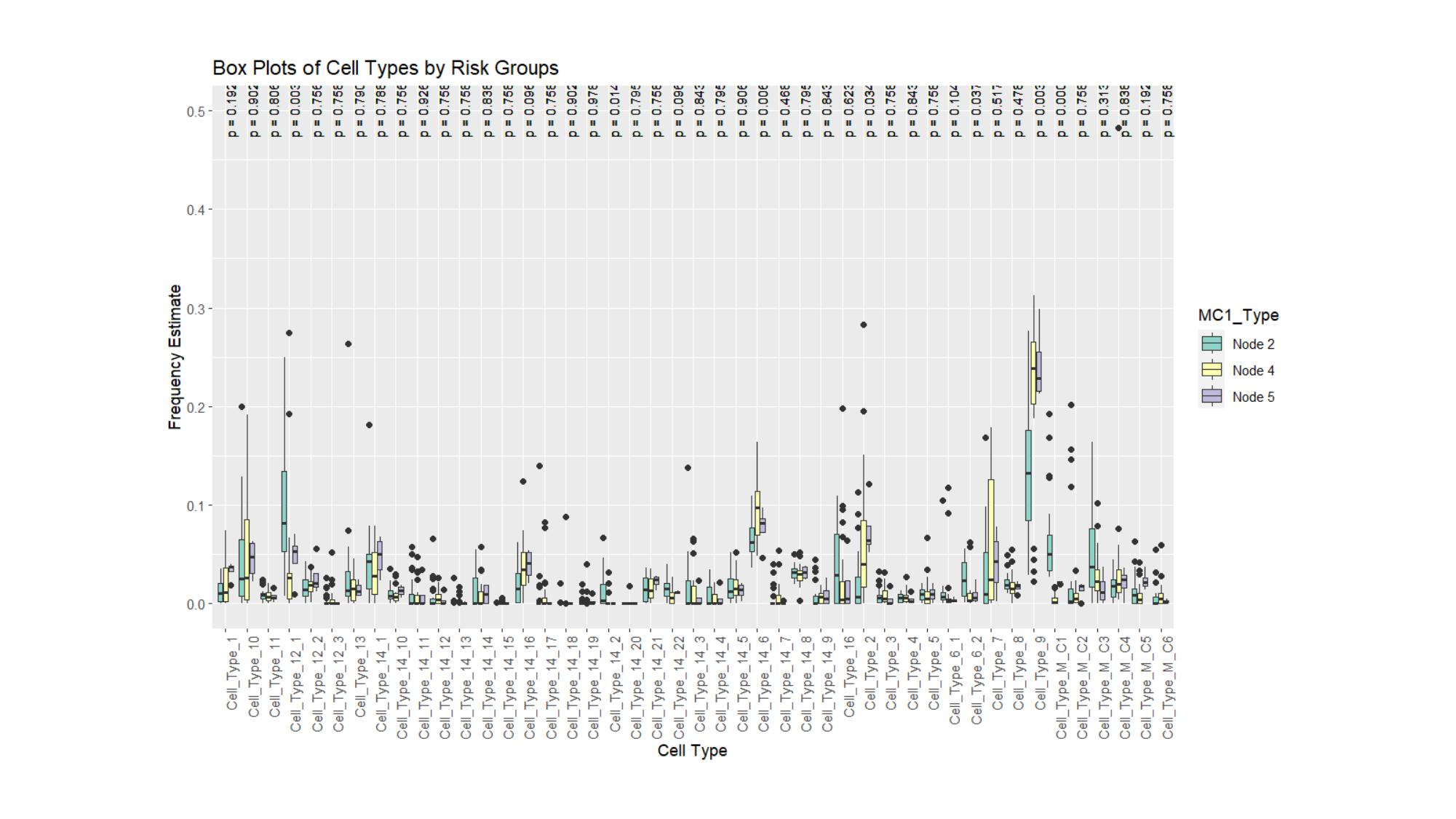

#### Slide 3
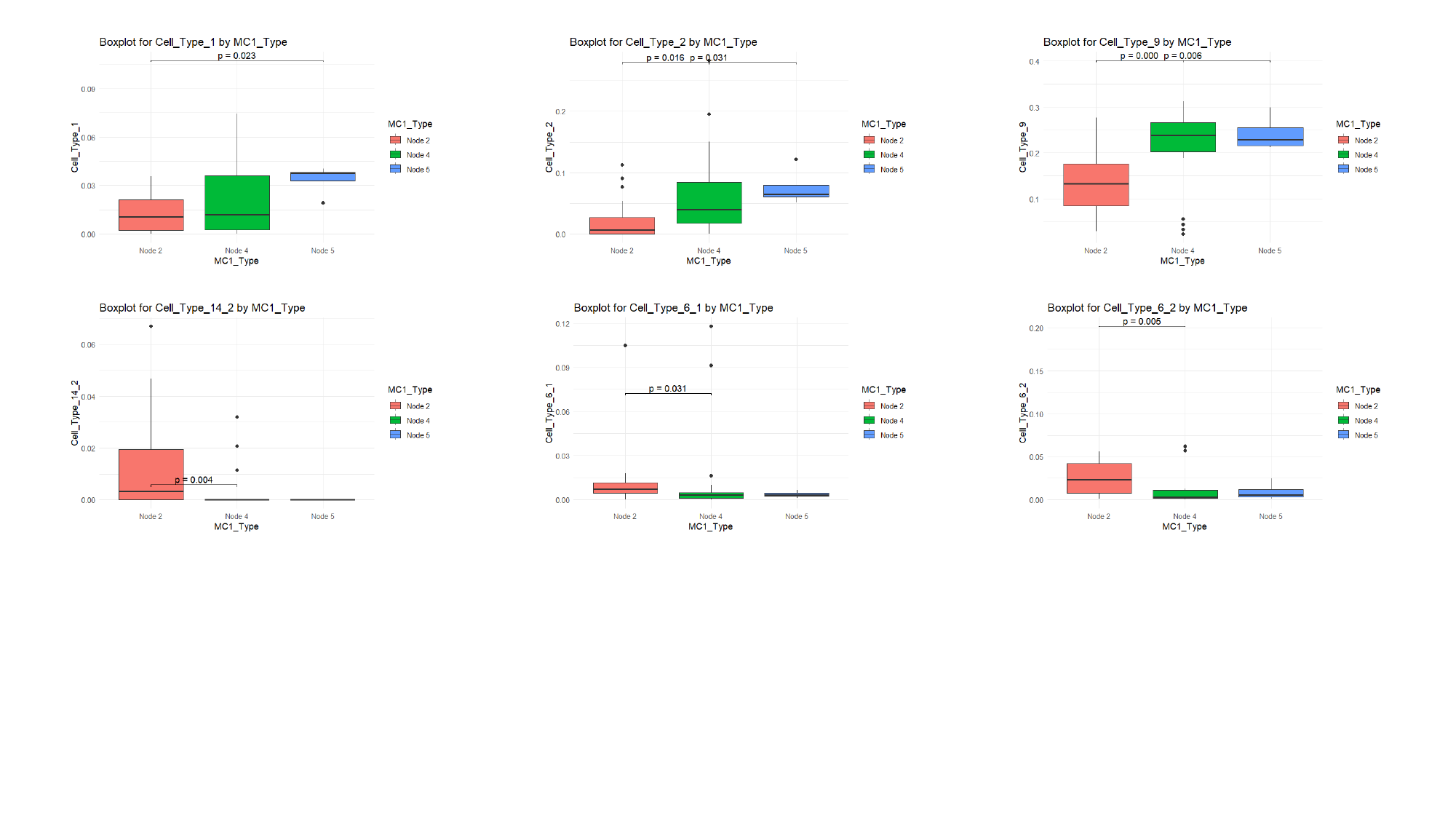

#### Slide 4
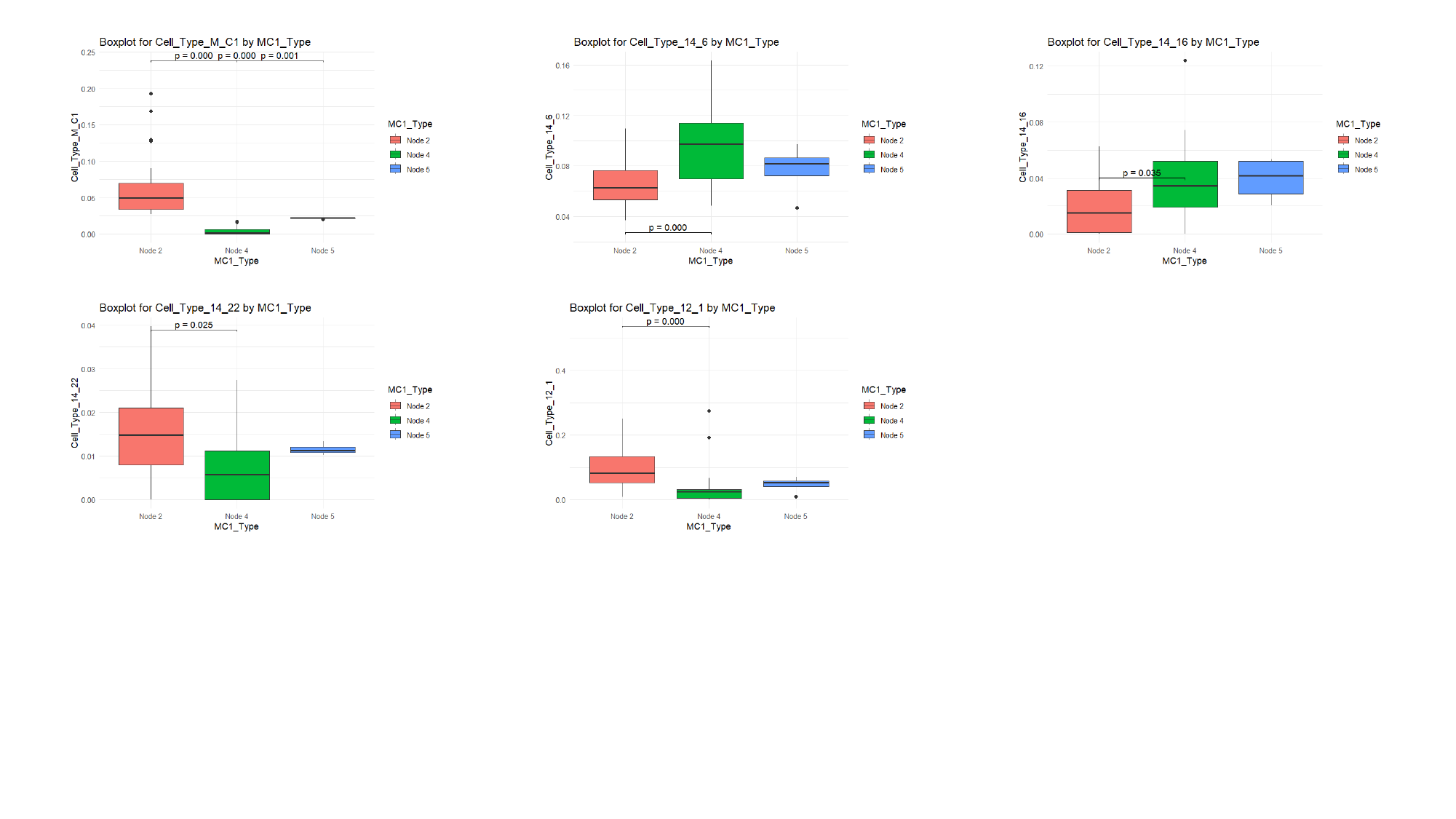

### Supplement 6

#### Slide 1
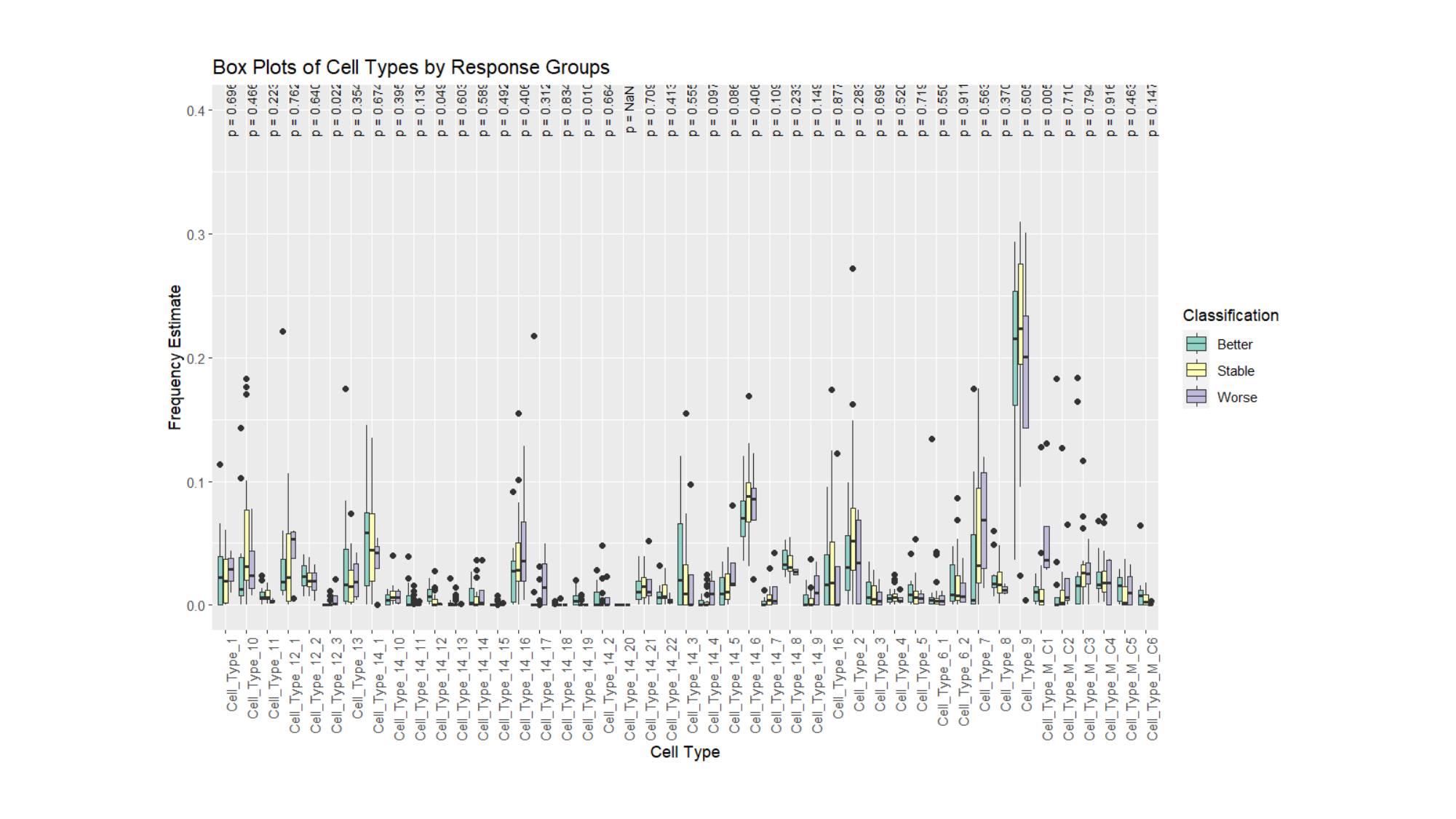

#### Slide 2
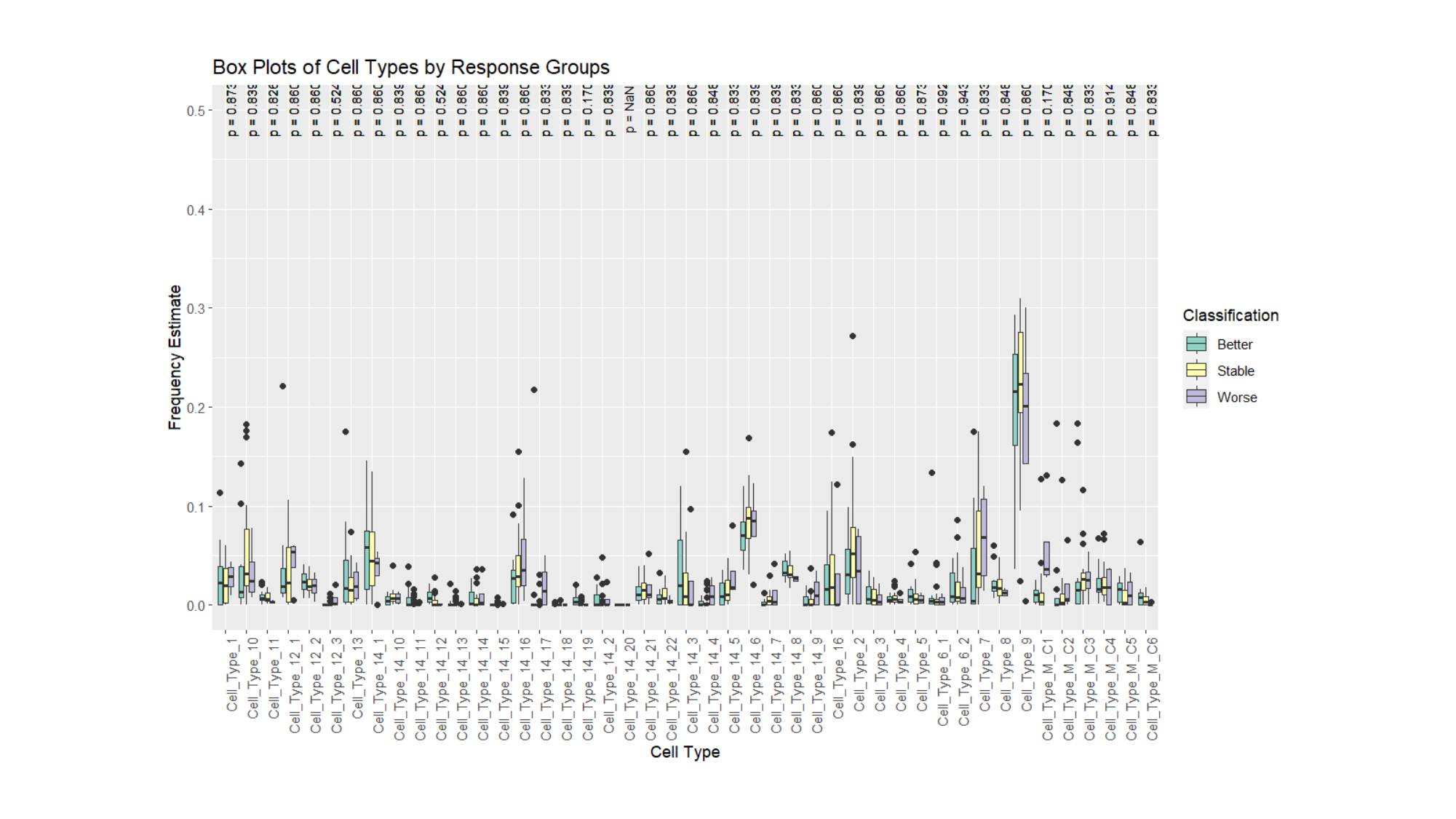

#### Slide 3
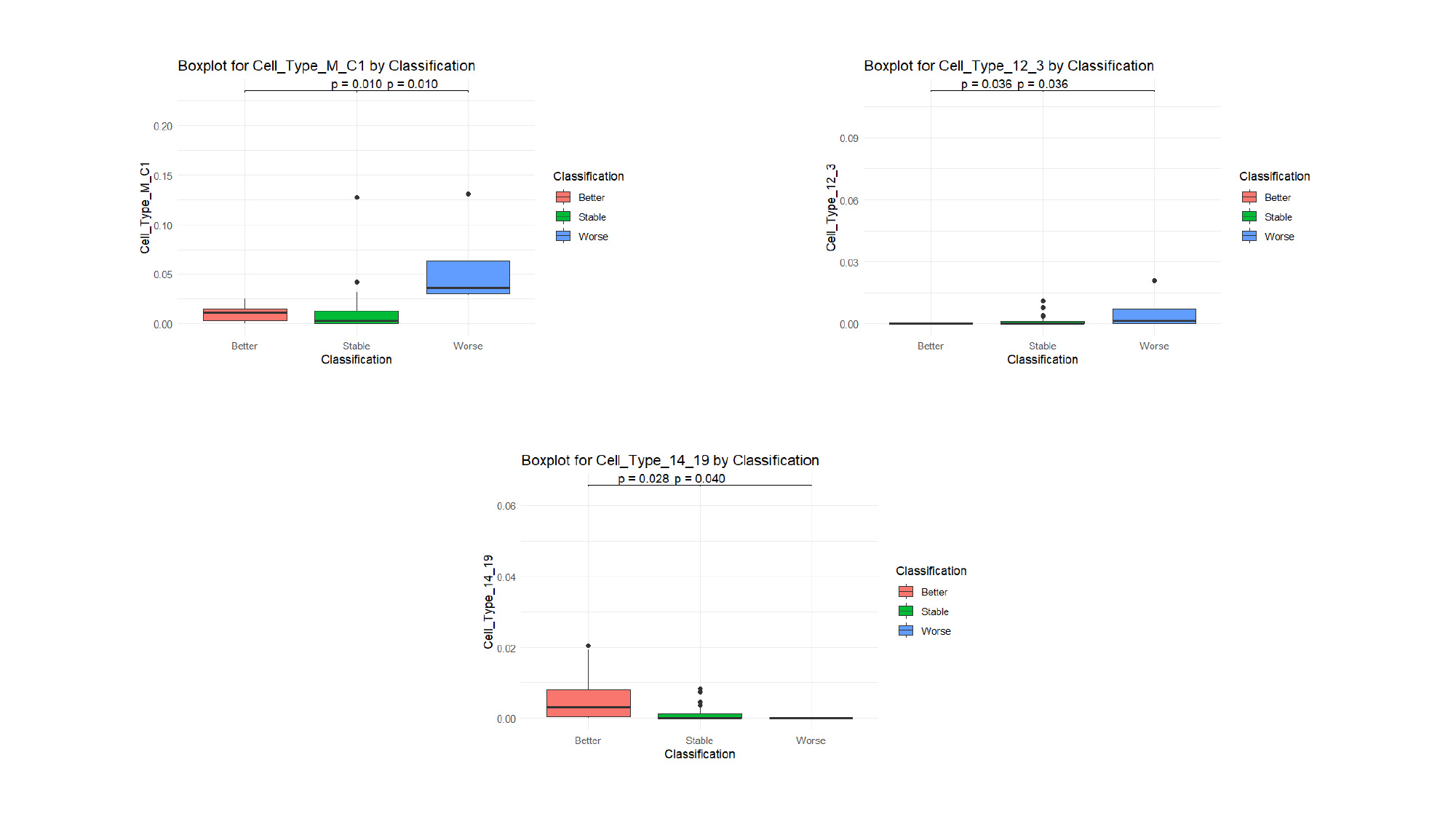

#### Slide 4
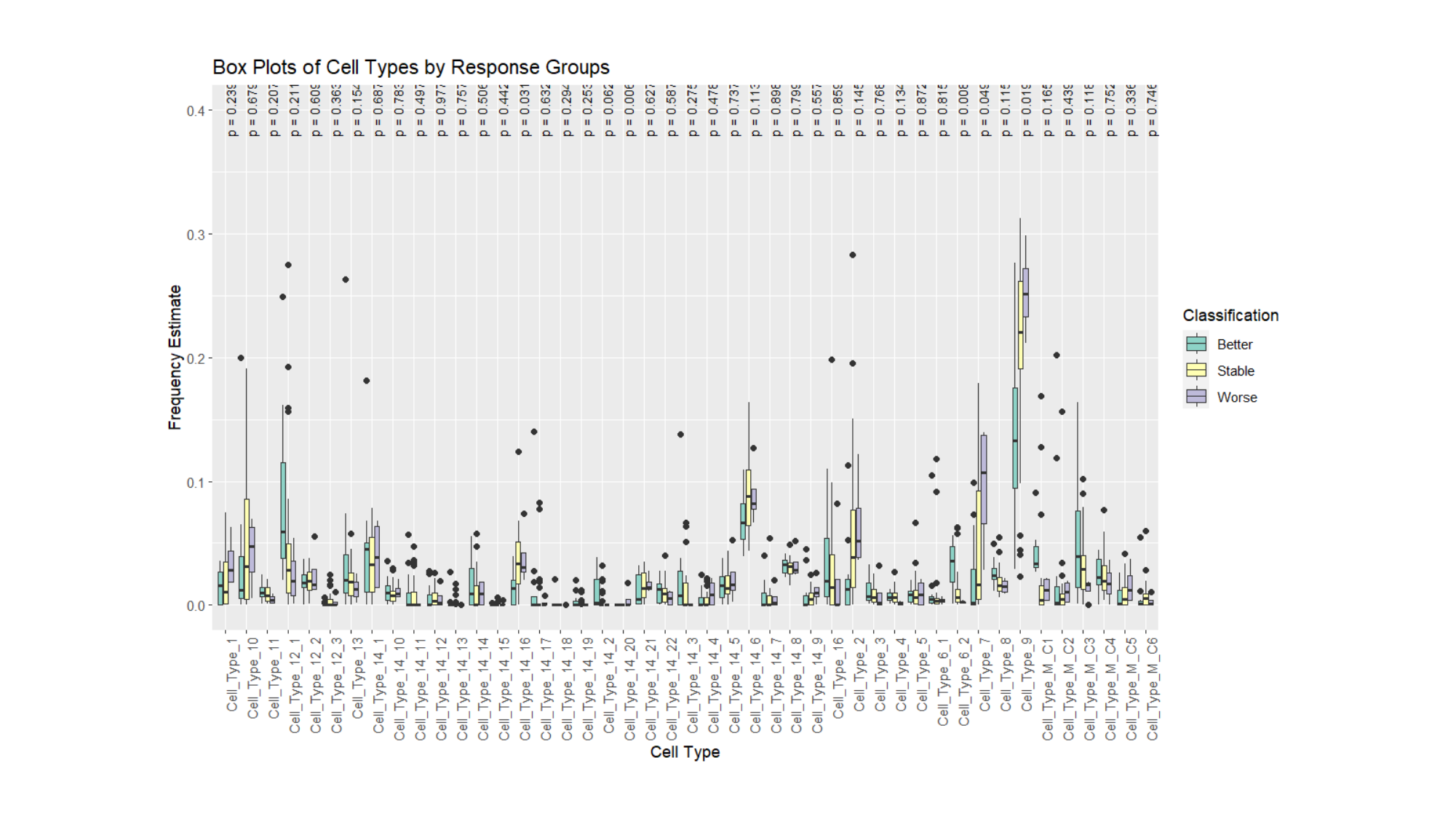

#### Slide 5
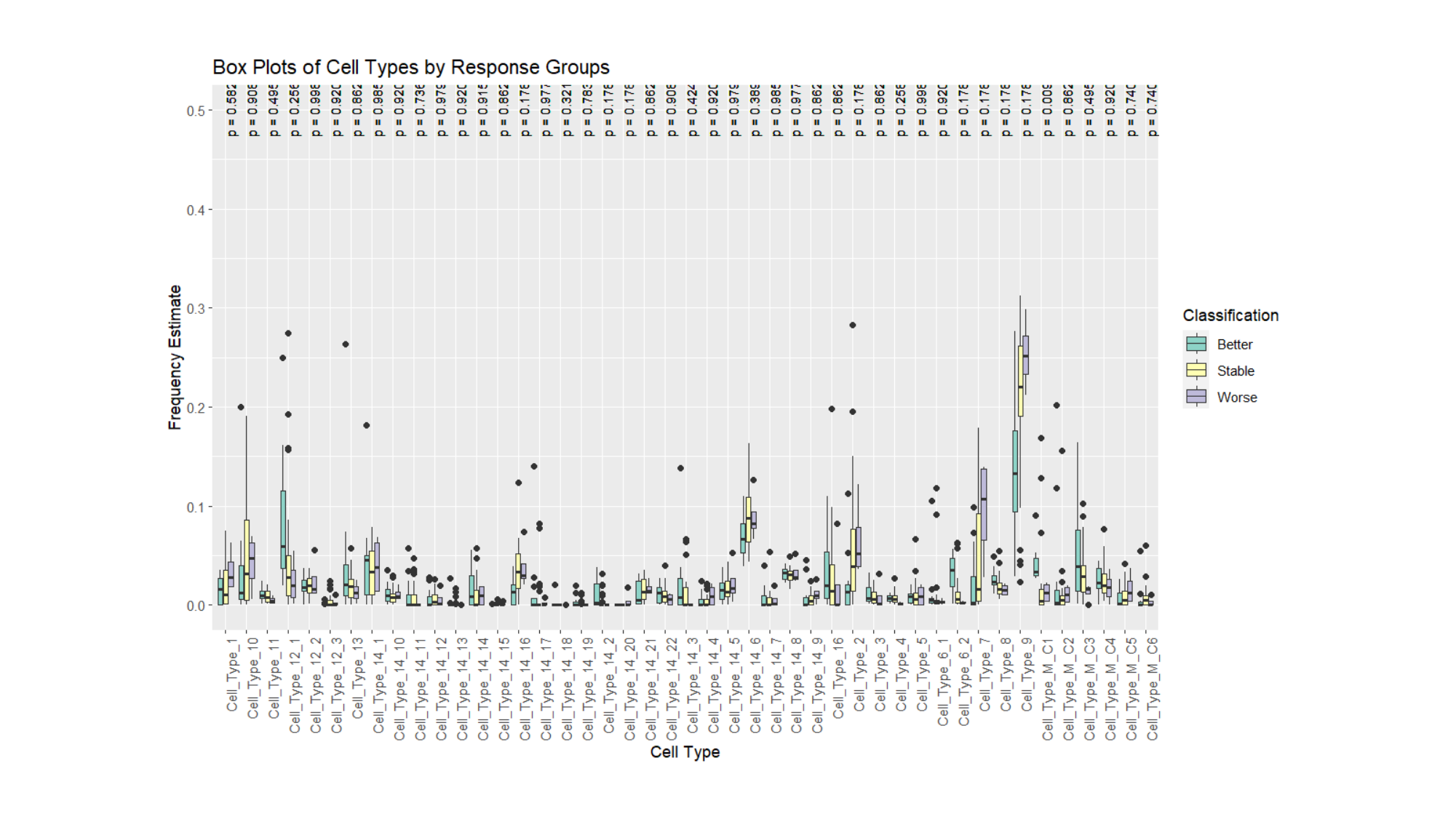

#### Slide 6
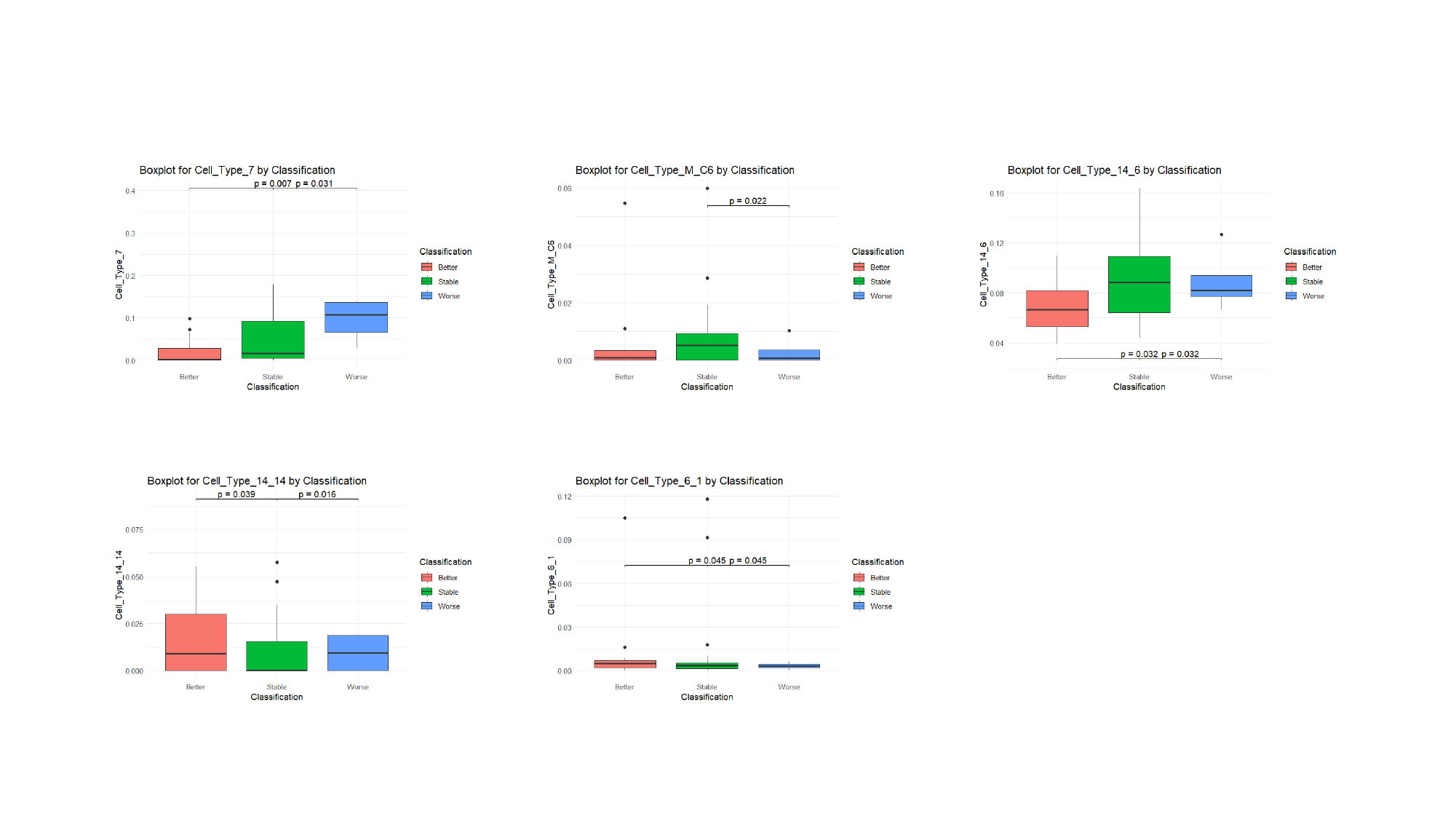

### Supplement 7

#### Slide 1
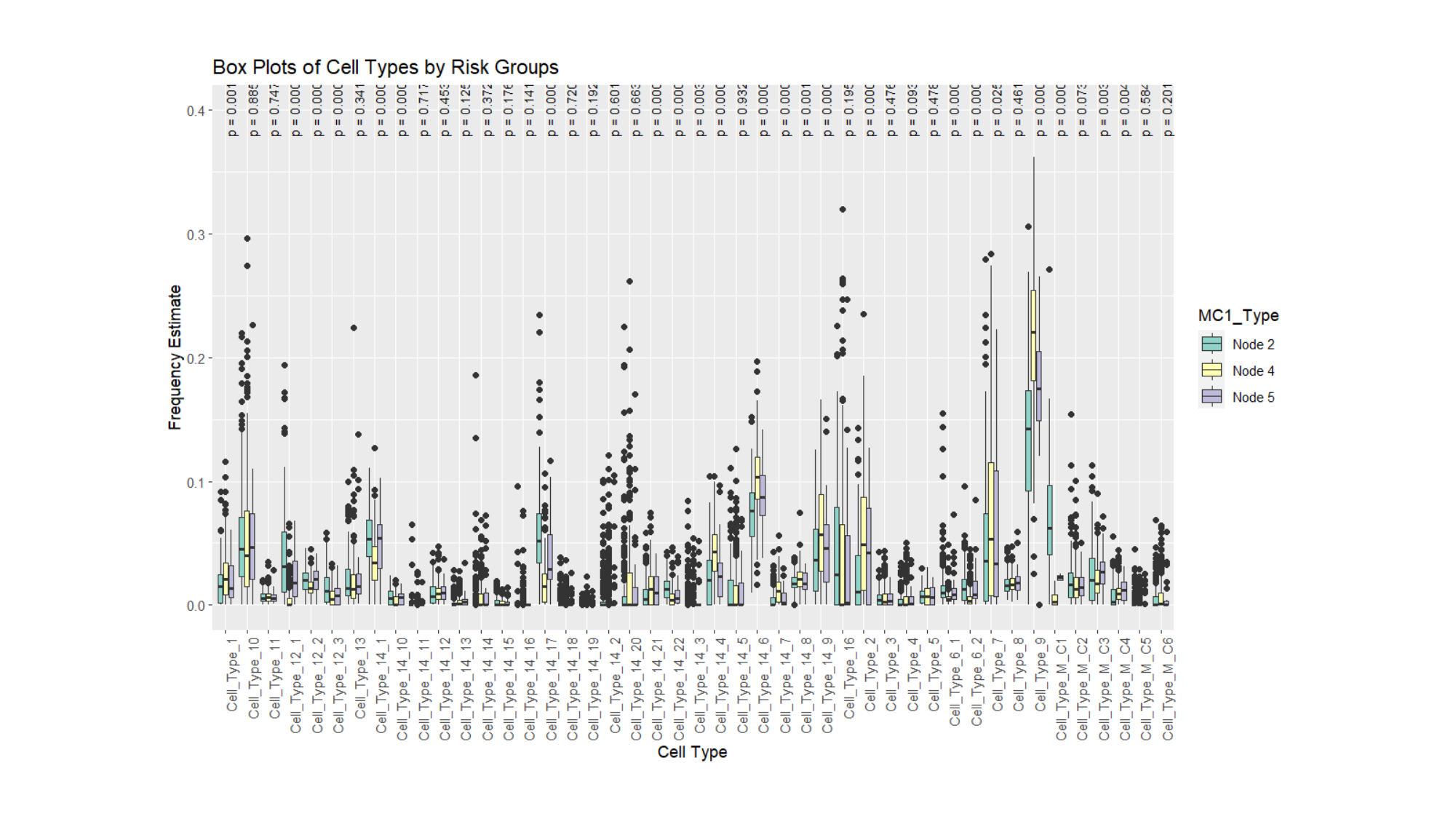

#### Slide 2
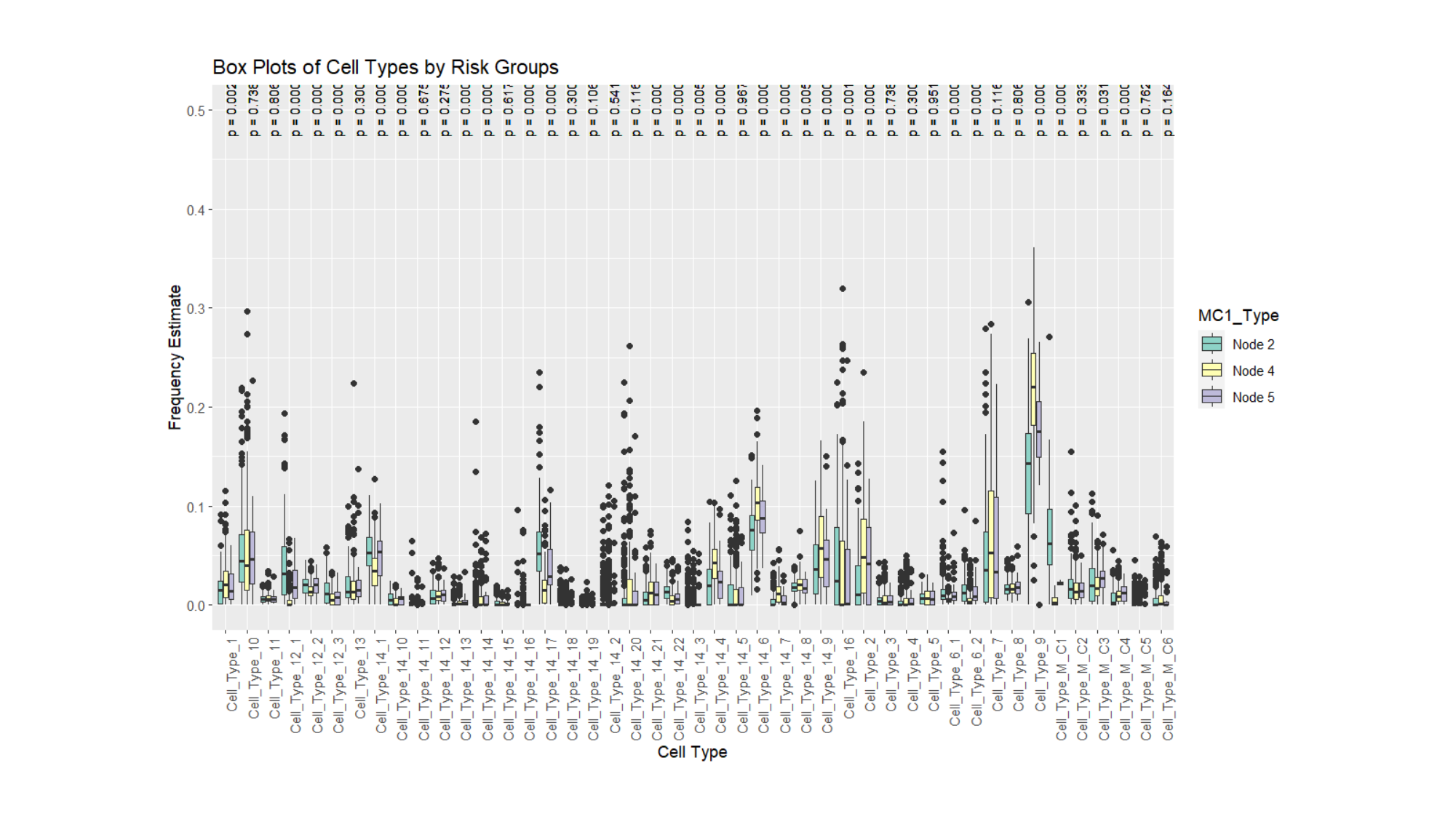

#### Slide 3
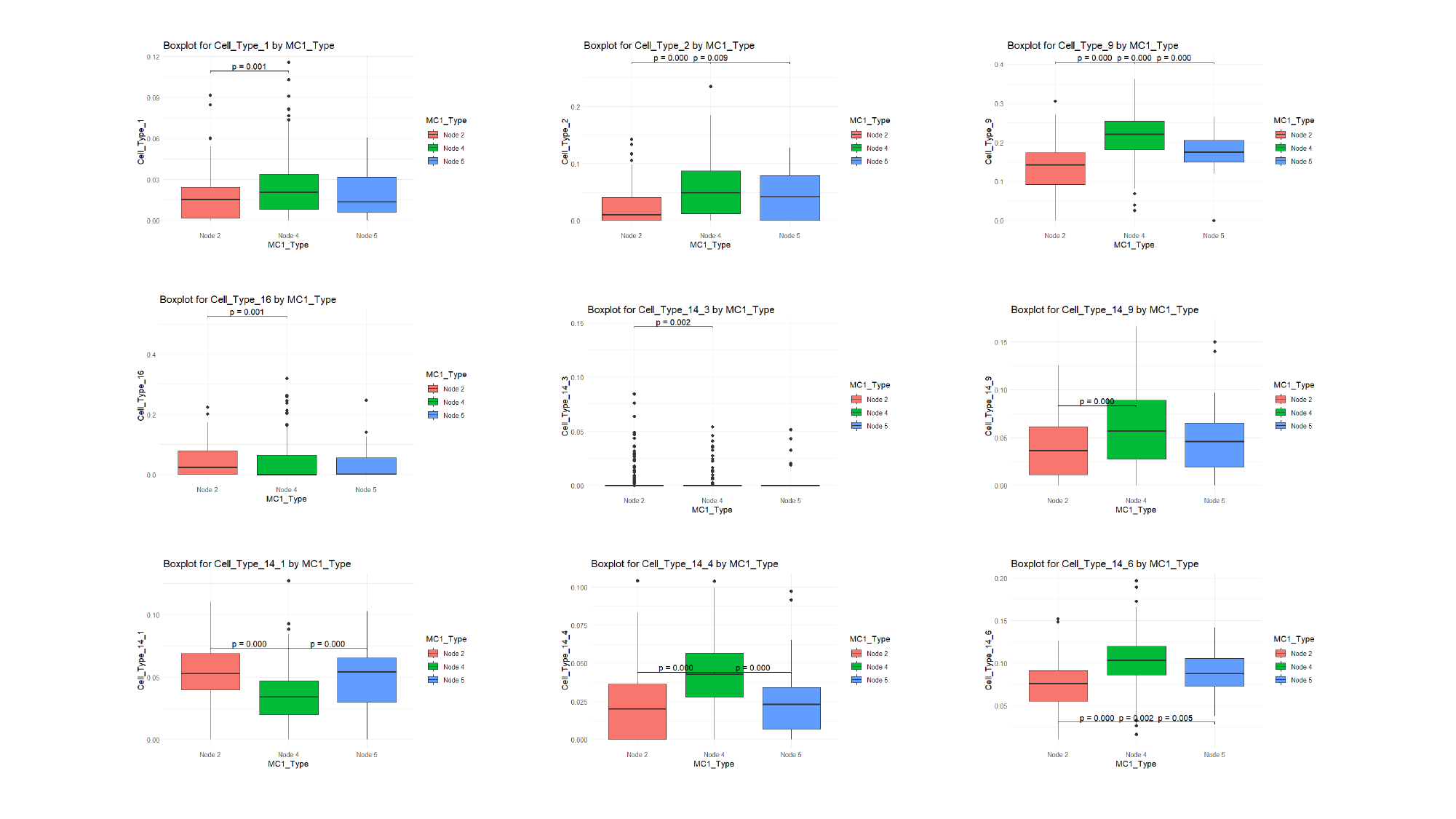

#### Slide 4
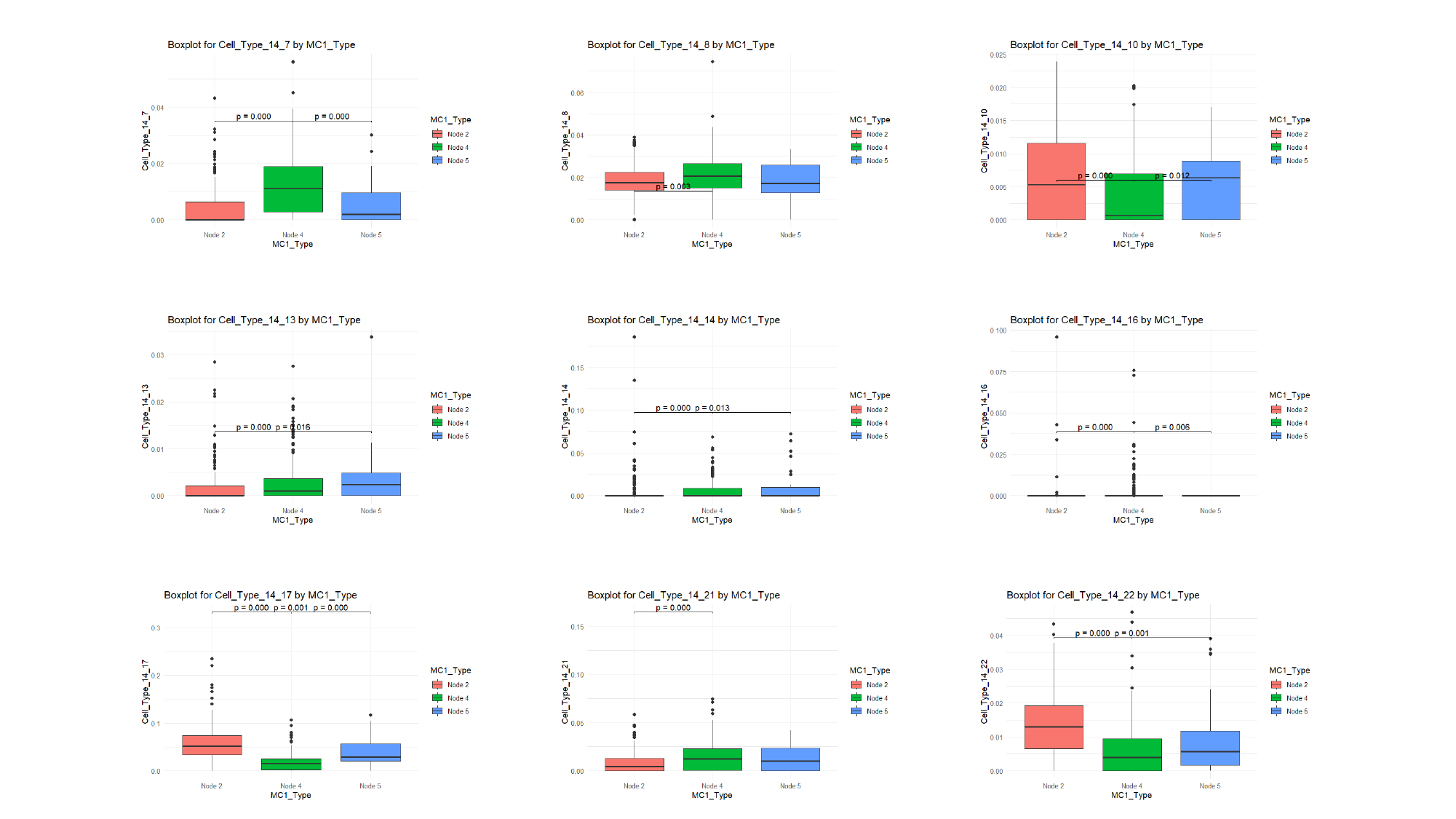

#### Slide 5
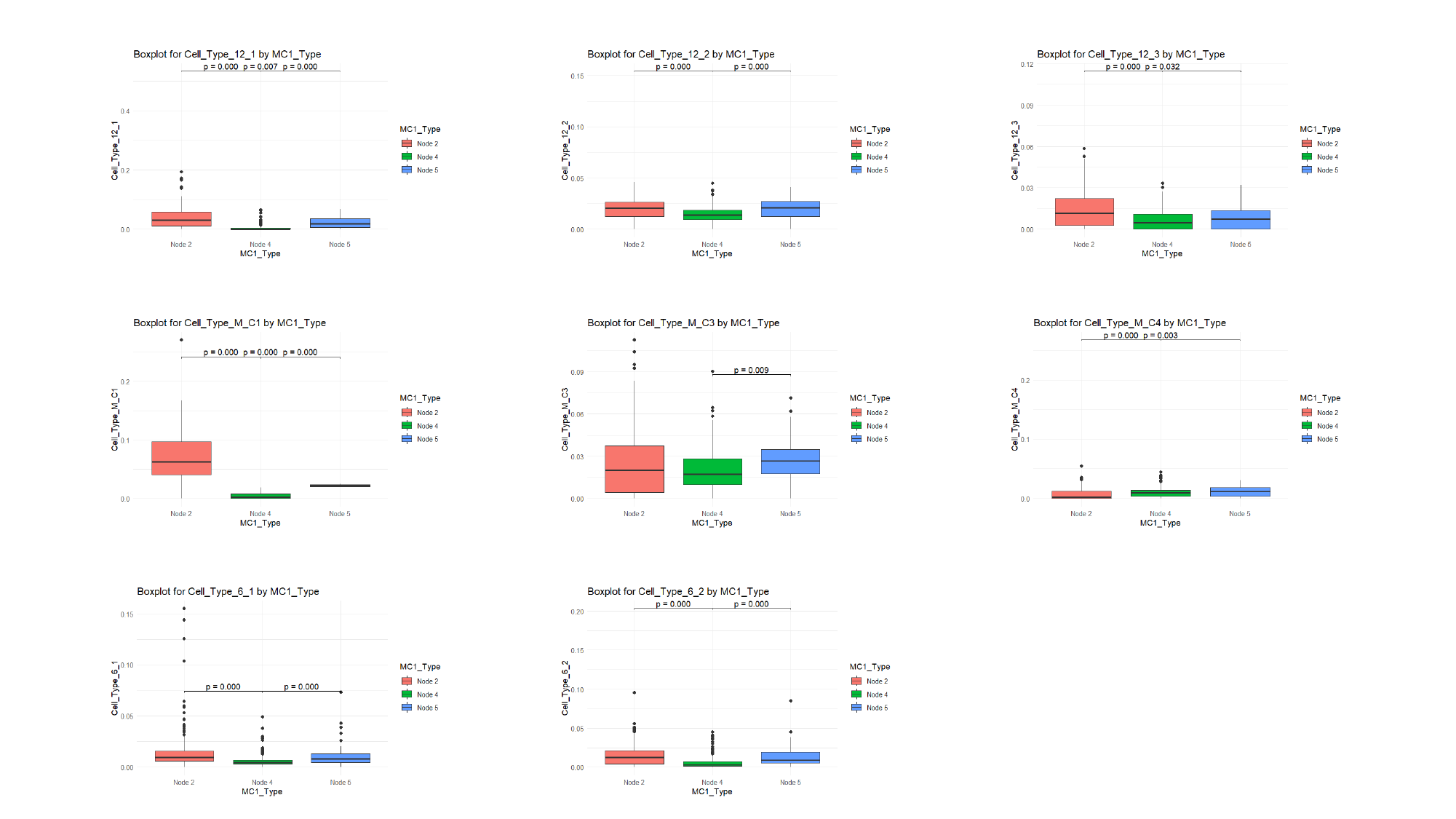
